# Genome-wide analysis of the class III peroxidase gene family in *Physcomitrium patens* and a search for clues to ancient class III peroxidase functions

**DOI:** 10.1101/2024.01.25.577222

**Authors:** Vincent P. M. Aparato, Fazle Rabbi, Taylor Madarash, Wyllie A. Brisbourne, Elizabeth I. Barker, Dae-Yeon Suh

## Abstract

Plant class III peroxidases (PRX) catalyze generation of reactive oxygen species and oxidation of various compounds including lignin precursors. PRXs function in cell wall metabolism, defense and stress responses. However, gene redundancy and catalytic versatility have impeded detailed functional characterization of *PRX* genes. The genome of the model moss *Physcomitrium patens* harbors a relatively small number (49) of *PRX* genes. Conserved architecture of four exons and three ‘001’ introns, found in some algal *PRX* genes and in the *PpPRX* family, suggests that this architecture predated divergence of the green algal and land plant lineages. The *PpPRX* family expanded mainly through whole-genome duplications. All duplicated pairs but one were under purifying selection and generally exhibited similar expression profiles. An expanded phylogenetic tree revealed a conserved land plant-wide clade that contained PRXs implicated in stress responses in non-lignifying cells, providing a clue to ancient functions of land plant PRXs. Functional clustering was not observed, suggesting convergent evolution of specific PRX functions (*e*.*g*., lignification) in different plant lineages. With its small complement of PRXs*, P. patens* may be useful for functional characterization of land plant PRXs. Several PpPRXs were proposed for further study, including *PpPRX34* and *PpPRX39* in the ancient land plant-wide clade.

## 1. Introduction

Peroxidases are a diverse group of enzymes that catalyze redox reactions in which substrates (e.g. monolignols) are oxidized while hydrogen peroxide (H_2_O_2_) or organic peroxides are reduced to water or corresponding alcohols. Based on the nature of the redox center, peroxidases are divided into heme or non-heme peroxidases (Passardi et al. 2007). Heme peroxidases are found in all kingdoms of life and comprise four superfamilies that evolved independently (Zámocký et al. 2015). Among them, the peroxidase-catalase superfamily is currently the largest superfamily and is composed of three structurally distinct families, which are termed class I, II and III peroxidases (Welinder 1992). Class I peroxidases are represented by ascorbate and cytochrome *c* peroxidases and their main function appears to be scavenging excess H_2_O_2_. Class II peroxidases include lignin-degrading fungal peroxidases (ligninases).

Secretory plant peroxidases such as horseradish peroxidases are categorized as class III peroxidases (PRXs). They are present as large multigene families in all land plants (embryophytes) and also in some streptophyte algae (Mbadinga Mbadinga et al. 2020). They are monomeric glycoproteins and are structurally characterized by four conserved disulfide bonds, two calcium ion binding sites and an active site with a heme prosthetic group (Veitch 2004). PRXs are secreted to the outside of the plant cell or to the vacuole and perform diverse tissue-specific functions. As examples, PRXs oxidize substrates in lignification, suberization, and auxin catabolism. They also generate or break down ROS (reactive oxygen species) in the processes of stress (e.g. pathogen) resistance, cell elongation and germination (Cosio and Dunand 2009; Shigeto and Tsutsumi 2016). A few examples of PRXs whose functions were studied by reverse genetics are: *Arabidopsis* PRX2, PRX25 and PRX71 in lignification (Shigeto et al. 2015), a pepper PRX (CaPO2) in ROS generation and disease resistance (Choi et al. 2007), and *Arabidopsis* PRX16 in seed germination (Linkies et al. 2010). Due to their diverse roles in plant growth and development as well as in defense responses, *PRX* families have been identified and characterized in model plants or major crops, including *Arabidopsis* (Tognolli et al. 2002; Welinder et al. 2002), rice (Passardi et al. 2004), poplar (Ren et al. 2014), maize (Wang et al. 2015), pear (Cao et al. 2016), switchgrass (Moural et al. 2017), wheat (Yan et al. 2019), cotton (Chen et al. 2022), grapevine (Xiao et al. 2020), potato (Yang et al. 2020), soybean (Aleem et al. 2022), and tobacco (Cheng et al. 2022).

Phylogenetic and gene network analyses have suggested that some PRXs likely emerged at the beginning of plant terrestrialization and that PRX families have expanded throughout plant evolution (Duroux and Welinder 2003; Oliveira et al. 2019). In turn, the antiquity and abundance of plant PRXs suggest that ancient plant PRXs may have played important roles in overcoming challenges encountered on land and, later, diversification of PRX functions may have helped plants to cope with increasing stresses during plant evolution. However, PRX families in vascular plants have a large number of members (e.g. 73 PRXs in *Arabidopsis* and 119 in maize) and their functional redundancy in addition to *in vitro* substrate promiscuity (Moural et al. 2017) make studying their contribution to plant terrestrialization and subsequent functional diversification difficult. Instead, PRX families of non-vascular plants with a smaller number of members can be advantageous in gaining insights to these questions (Francoz et al. 2015).

*Physcomitrium patens* has emerged as a model moss (Rensing et al. 2020). Owing to the ease of transformation and vegetative propagation of transformed lines, gene knockout phenotypes can be readily screened (Hohe et al. 2004). Its genome sequence is well-annotated, and large-scale tissue-specific gene expression data are available (Fernandez-Pozo et al. 2020). *P. patens* is also amenable to chemical rescue studies because its single cell layered tissues can uptake chemicals through the entire surface (Strotbek et al. 2013). In line with its simple morphology and physiology, the number of *PRX* genes in *P. patens* is roughly half that in vascular plants. These characteristics make *P. patens* an excellent system for studying the functions and evolution of *PRX* genes with reverse genetics and chemical approaches. We are interested in how certain ancient PRXs contributed to the early evolution of land plants, especially in the metabolism of protective biopolymers (e.g. sporopollenin, lignin-like materials) and in spore development. In this study, we performed, for the first time for any bryophyte, a genome-wide analysis of the *PRX* gene family in *P. patens*. We identified forty-nine *PRX* genes and two pseudogenes and analyzed their deduced amino acid sequences, phylogeny, gene duplication events and expression profiles. We also constructed large scale phylogenetic trees in search of PRXs that are highly conserved in land plants and possibly possess ancient functions.

## 2. Methods

### 2.1. Identification and sequence analysis of class III peroxidase genes in *P. patens*

*P. patens* PRX sequences collated in Lehtonen et al. (2009) and in RedoxiBase (https://peroxibase.toulouse.inra.fr) (Savelli et al. 2019) were used for tBLASTn searches of the *P. patens* v3.3 genome in Phytozome 13 (https://phytozome-next.jgi.doe.gov). Protein sequences of the candidate genes were further analyzed for the conserved amino acid residues (Veitch 2004) and named from PpPRX1 to PpPRX51 according to their chromosomal locations.

Subcellular localization was predicted using DeepLoc 2.0 (https://services.healthtech.dtu.dk/services/DeepLoc-2.0/) (Thumuluri et al. 2022). Transmembrane domains were predicted by Phobius, a combined transmembrane topology and signal peptide predictor (https://phobius.sbc.su.se/) (Käll et al. 2004). Multiple sequence alignment was produced using the MAFFT L-INS-I method (https://mafft.cbrc.jp/alignment/server/).

### 2.2. Phylogenetic analysis

A Maximum Likelihood (ML) phylogenetic tree of the PpPRX family was reconstructed from amino acid sequences encoded by 49 *PpPRX* genes and 2 pseudogenes. The sequences were aligned using the MAFFT L-INS-I method and an ML tree was built in MEGA X (Kumar et al. 2018) under the JTT substitution model. The initial tree was created using the default tree inference options. Support for the tree was measured using 1,000 bootstrap replicates. The bootstrap consensus tree was then formatted using iTOL (Letunic and Bork 2019).

An ML tree of all PpPRXs and PRXs selected from eight model plants, each representing a major embryophyte lineage, was also reconstructed to find any embryophyte-spanning clades. Amino acid sequences of PRXs from *Anthoceros punctatus*, *Marchantia polymorpha*, *Selaginella moellendorffii*, *Ginkgo biloba*, *Amborella trichopoda*, *Oryza sativa* and *Arabidopsis thaliana* were retrieved from RedOxibase, and *Ceratopteris richardii* PRX sequences were obtained from Phytozome 13. Then, eight single-species ML trees were reconstructed and, from each tree, a set of phylogenetically distinct PRXs—all singletons and one from each multi-tip clade—was selected. For example, twenty-two *A. punctatus* PRXs (ApPRXs)—seven singletons and one each from fifteen multi-tip clades—were selected from an ML tree of fifty-four ApPRXs. Then, several multi-species ML trees were reconstructed to remove species-specific clades. For example, an ML tree was reconstructed from ApPRXs and *A. thaliana* PRXs (AtPRXs) and any PRX either in ApPRX-specific or in AtPRX-specific clade was removed. The selection and pruning of PRXs were repeated until only PRXs that were in multi-species clades remained. Lastly, 49 PpPRXs and 32 non-redundant PRXs with reported functions (Table S1) were added to give a total of 247 PRXs. An ML tree was reconstructed in MEGA X as described above except that the sequences were aligned using the MAFFT FFT-NS-i method and the WAG substitution model with the shape parameter of the gamma distribution, a proportion of invariable sites and empirical amino acid frequencies was used (WAG+G+I+F). The number of discrete gamma categories was set to 4. The tree was rooted with an ascorbate peroxidase sequence from the streptophyte alga *Spirogloea musicola* CCAC0214. Support for the tree was measured using 1,000 bootstrap replicates.

### 2.3. Gene duplication analysis

Gene pairs were investigated for duplication events when two genes shared ≥70% protein sequence identity, or shared <70% sequence identity and had one or more companion gene pairs. Duplicated gene pairs were investigated for their phylogenetic relationship, Ks/Ka ratio, gene architecture, expression profile, and companion pairs in 50 kb flanking regions. In the lineage leading to *P. patens*, two whole-genome duplications (WGD) occurred (Lang et al. 2018). In WGD1, seven ancestral (anc7) chromosomes were duplicated and subsequent loss of a chromosome resulted in thirteen (anc13) chromosomes. Following WGD2, fissions and fusions resulted in the extant twenty-seven chromosomes. A duplicated pair was assumed to be produced during WGD1 if the genes were located in two chromosomes that descended from the same anc7 chromosome but from different anc13 chromosomes. Similarly, a duplicated pair was assumed to be produced during WGD2 if the genes were present on two chromosomes that were descendants of the same anc13 chromosome. Two nearby duplicated genes were defined as tandemly duplicated (TD) genes when they were less than 100 kb apart and there were fewer than three intervening genes, or segmentally duplicated (SD) genes otherwise.

Ka and Ks values were obtained by first performing pairwise alignment of the coding region nucleotide sequences using MAFFT in RevTrans (Wernersson and Pedersen 2003). Each alignment was then imported into DnaSP v6 (Rozas et al 2017) to calculate asynonymous (Ka) and synonymous substitutions (Ks).

### 2.4 Gene expression analysis

*P. patens* developmental stage expression data were collated from the database PEATMoss (Fernandez-Pozo et al. 2020). To determine the number of clusters, wss and silhouette methods of Kmeans clustering were used where nstart was set at 25. Hierarchical clustering was calculated using the heatmap.2 package, employing the WardD2 and the Pearson correlation methods through R version 4.2.2. Gene and tissue dendrograms were generated using data obtained from Ortiz-Ramirez et al. (2016). The heatmap was generated from the same data using log2 transformed expression data in Microsoft Excel.

## 3. Results

### 3.1. Identification and sequence analysis of PpPRXs

A total of 49 *PRX* genes and 2 pseudogenes were identified in the *P. patens* genome (Table 1). While most are single copy genes, four genes have multiple copies and *PpPRX50* has four copies. *PpPRX49* and *PpPRX50* are reported here for the first time. To avoid mis-annotation of the other heme peroxidases that share parts of common PRX sequences (Mathé and Dunand 2021), we used the following criteria:

1. Two signature sequences, RLxF<u>H</u>DC_2_xxxGC_3_DxS and LxGx<u>H</u>xxGxxxC_6_, harboring the distal and proximal <u>H</u>is residues, respectively, of the heme (Fig. S1, Passardi et al. 2004).
2. Eight conserved Cys (Cys1 to Cys8) that form four disulfide bonds.
3. Conserved residues at the two calcium binding sites, including Asp71, Asp78, and Ser80 for the distal site, and Ser195, Asp239, Thr242, and Asp247 for the proximal site (numbering of PpPRX1, Fig. S1, Veitch et al. 2004).
4. Conserved gene architecture (intron locations and phases).
5. Signal peptide.

**Table 1.**
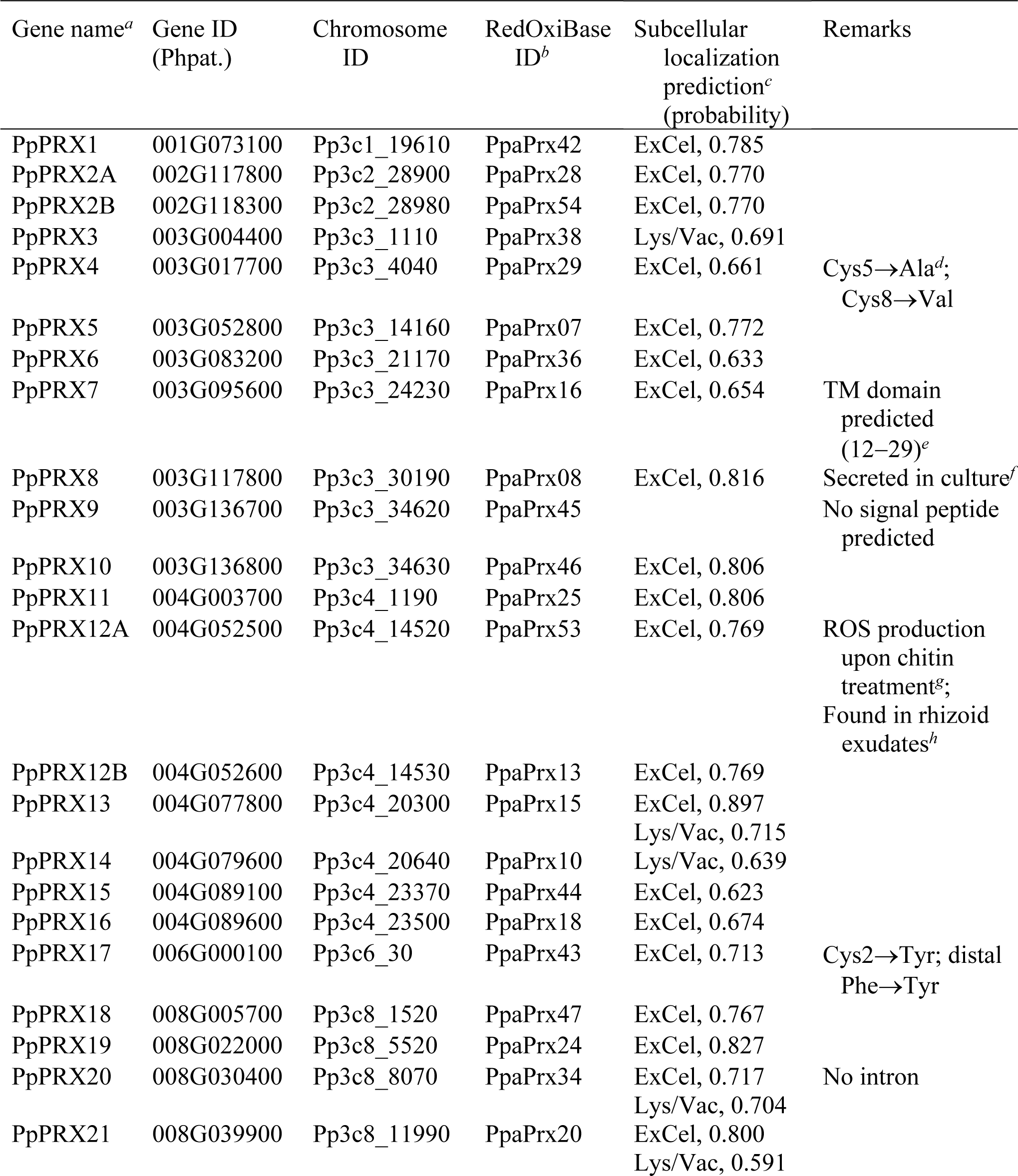
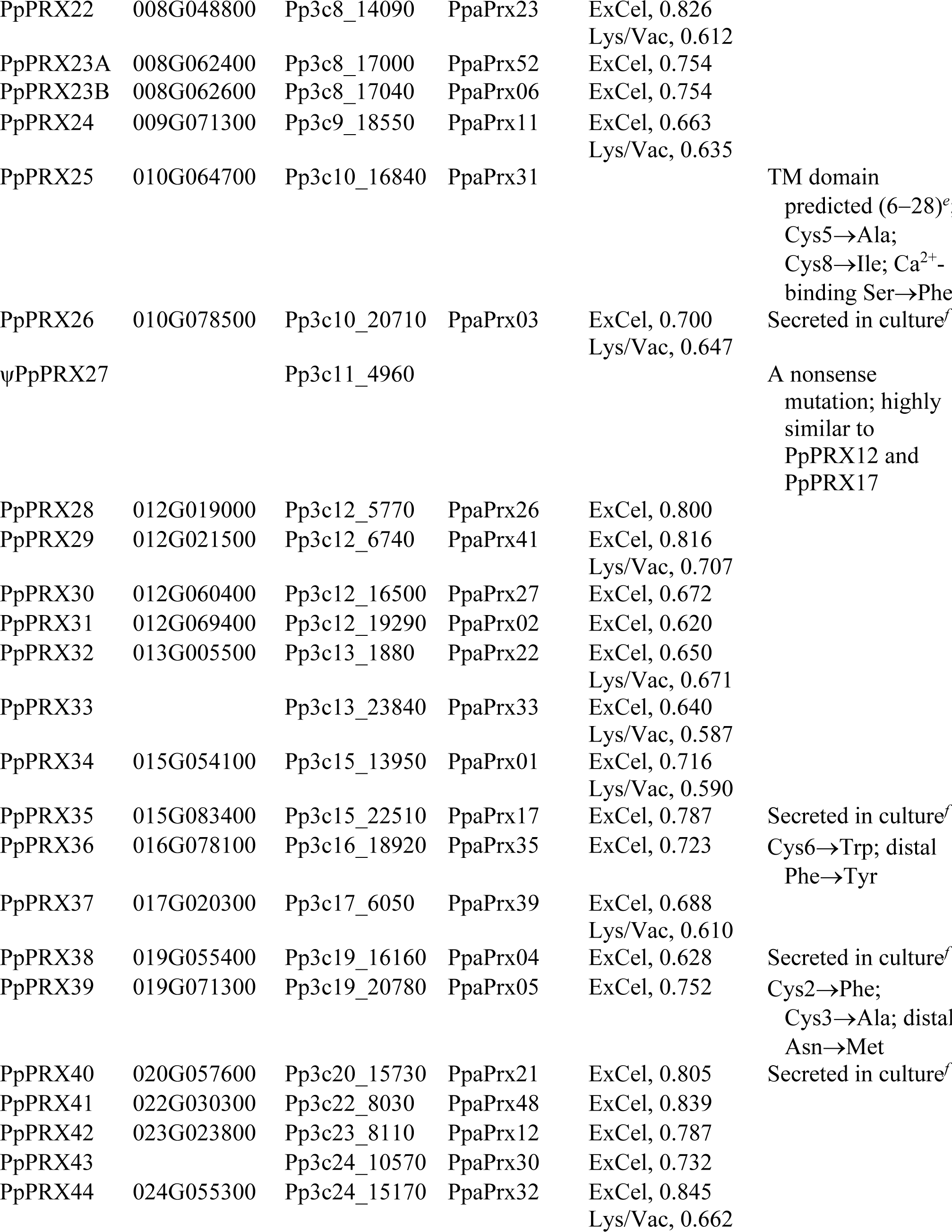
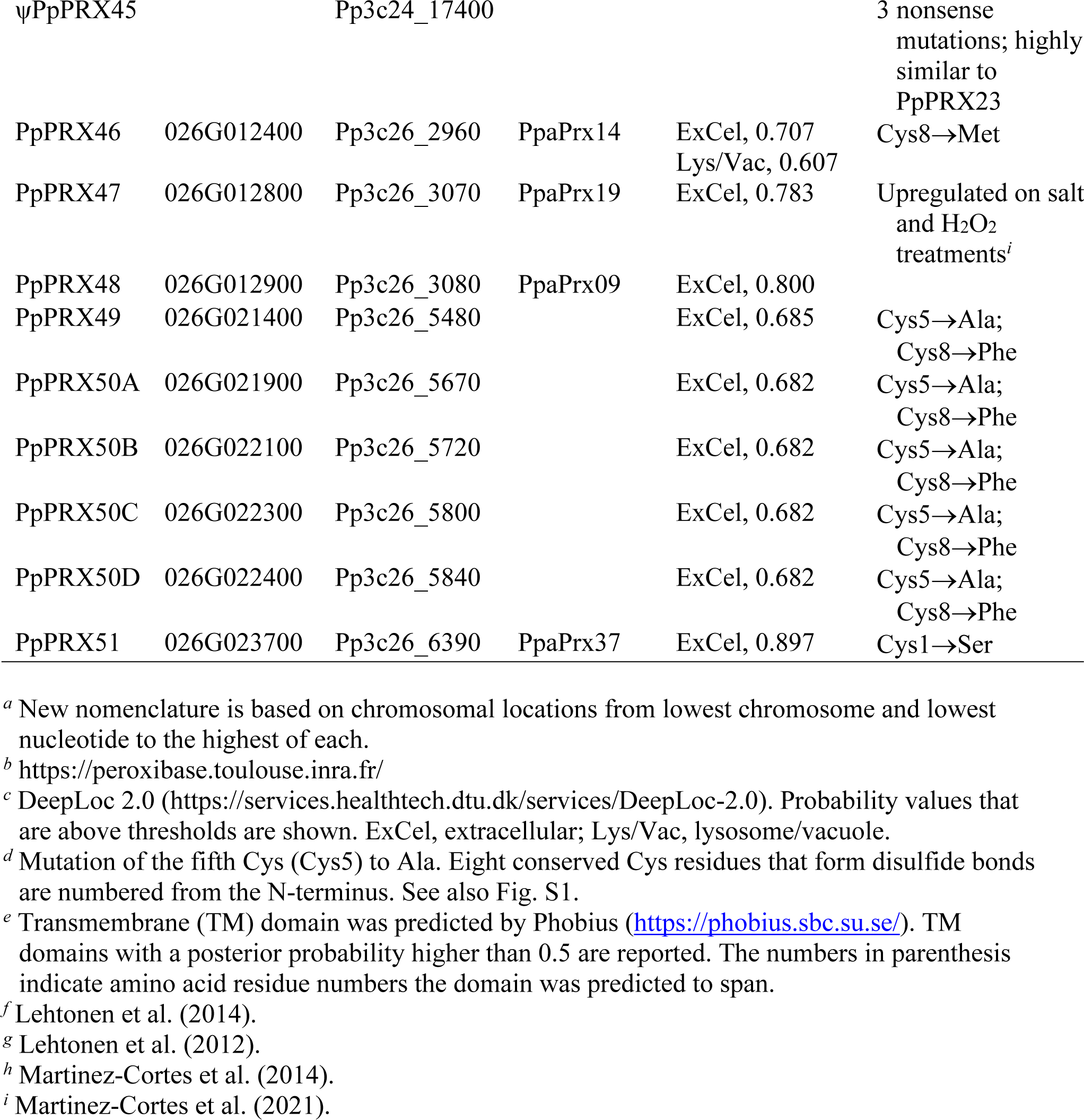
*P*. *patens PRX* genes and pseudogenes. Fifty-five *P. patens* genes and two pseudogenes are listed along with their identifiers, predicted subcellular localizations and additional notes, including reported functions, predicted transmembrane domains, non-conservative amino acid substitutions and other notable features.

A few proteins were tentatively assigned to be PRXs, although they did not meet all the criteria. For examples, PpPRX9 and PpPRX25 did not contain a predicted signal peptide, while PpPRX7 and PpPRX25 were predicted to be membrane-bound (Lüthje and Martinez-Cortes 2018). Several proteins including PpPRX4 and PpPRX17 lacked one or two of the eight conserved Cys residues (Table 1). When two Cys residues were mutated, both were always a disulfide forming pair, as Cys2 and Cys3 in PpPRX39, and Cys5 and Cys8 in PpPRX4, PpPRX25, PpPRX49 and PpPRX50. Hence, no PpPRX potentially lacking more than one disulfide bond was identified in this study. In an earlier study, a small fraction (<5%) of plant PRXs were predicted to lack one or two disulfide bonds (Mbadinga Mbadinga et al. 2020). Conserved residues in PpPRXs and in several PRXs from streptophyte algae and tracheophytes are highlighted in Fig. S1.

There are four multicopy genes. *PpPRX2*, *PpPRX12* and *PpPRX23* have 2 copies each, while four copies of *PpPRX50* are found. The lengths of untranslated regions that are identical in nucleotide sequence among gene copies are generally in the range of 700 to 2300 bp.

### 3.2. Phylogenetic analysis of the PpPRX family

The rooted ML tree of PpPRX sequences comprised seven major clades, A to F, and clades C and F were further divided into multiple subclades (Fig. 1). Except for the singleton clade C1 (PpPRX41), all nodes were supported by bootstrap values higher than 50%. The seven major clades were hierarchical: clade A of PpPRX34 and PpPRX39 was sister to the rest of the family, while clade B was sister to clades C−F. Clade D consisted of a single gene, PRX30, sister to clades E and F.

**Fig. 1.**
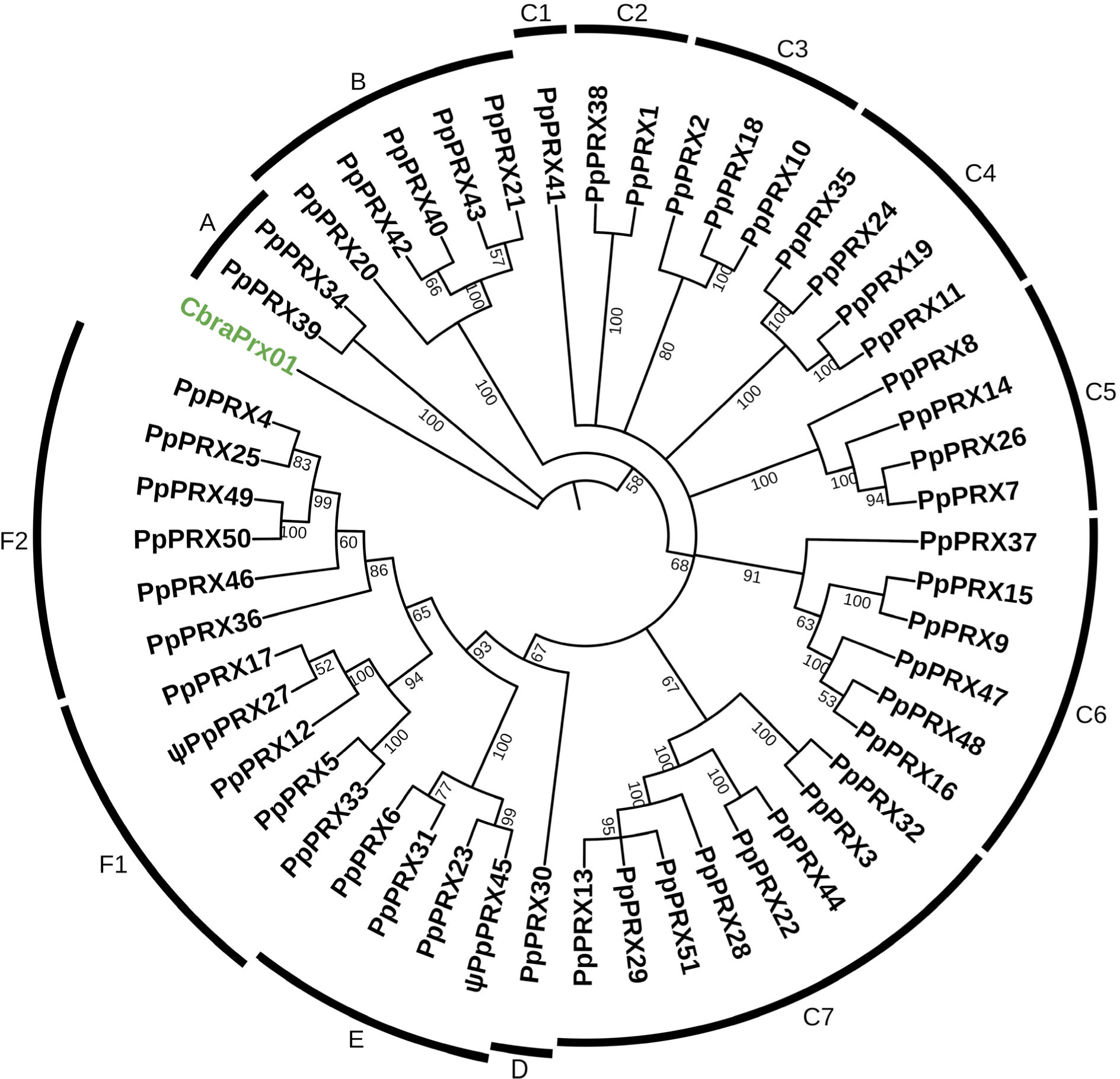
Maximum Likelihood phylogenetic tree of *P. patens* PRX family. A Maximum-Likelihood tree was constructed in MEGA X with the JTT substitution model and rooted with an algal PRX sequence from *Chara braunii* (CbraPrx01, GenBank accession id, GBG90290.1). Numerical values at nodes indicate bootstrap support above 50% from 1000 bootstrap replicates. Clades of the same phylogenetic level are labeled from A to F and sister clades within each level are labeled numerically.

### 3.3. Gene architecture

Of 49 *PpPRX* genes, 18 genes have three introns at conserved positions (Figs. 2, S1). Introns 1 and 2 are phase 0 introns, while intron 3 is a phase 1 intron. As shown in Fig. 2, these ‘001’ introns are not only highly conserved in plant *PRX* genes (e.g., horseradish *PRX C1*), but also are found in some streptophyte algal genes (e.g., *Klebsormidium nitens PRX* and *Spirogyra* sp. *PRX03*). Intron 1 is positioned between Cys2 and Cys3, downstream of the distal His residue, while intron 3 is positioned upstream of the proximal His residue (Fig. S1). The other 27 *PpPRX* genes experienced at least one intron loss, whereas four genes (*PRX11*, *PRX22*, *PRX44*, *PRX25*) underwent intron loss and gain events (Fig. 2).

**Fig. 2.**
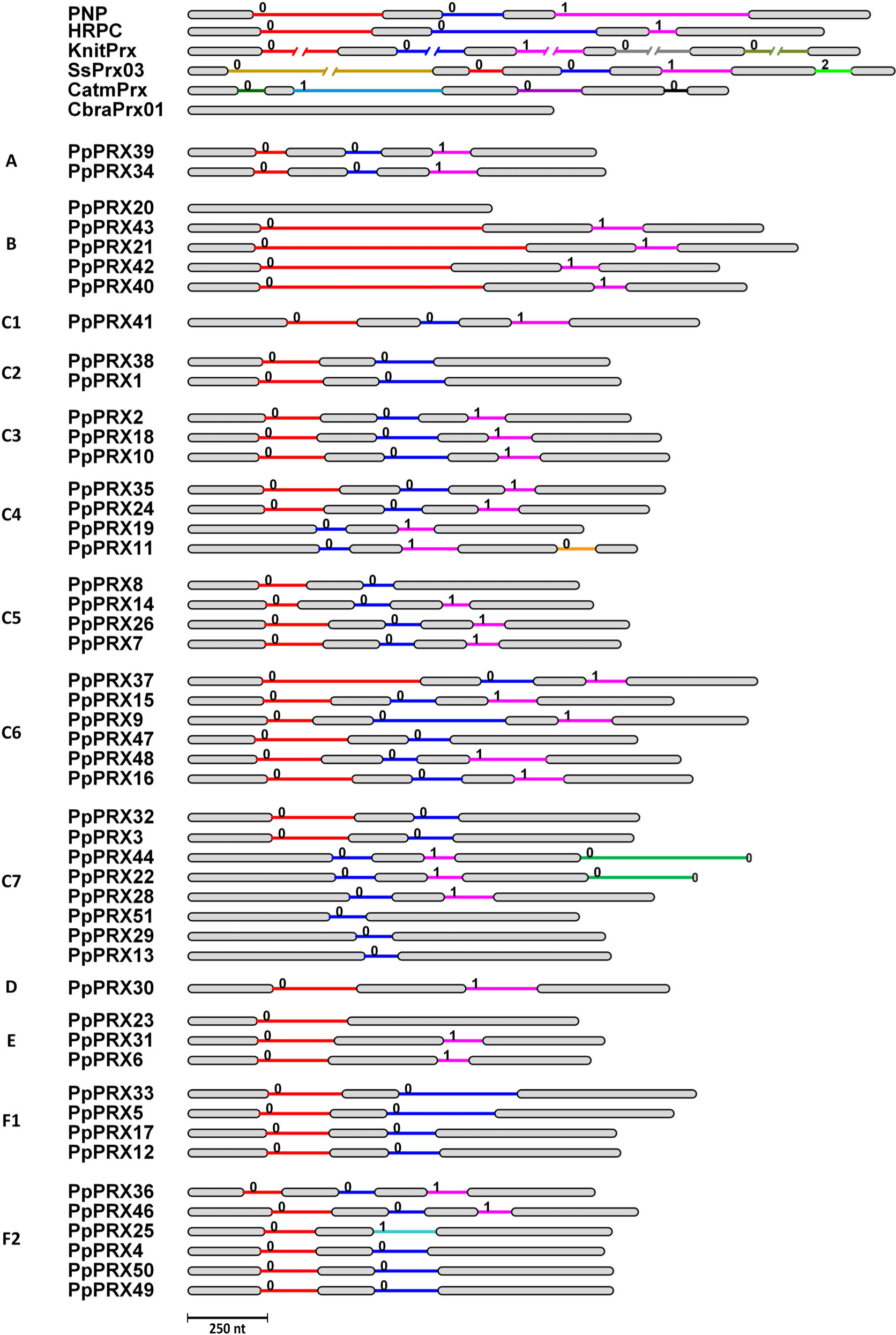
Architecture of *P*. *patens PRX* genes and conservation of their intron positions and phases. Each gene is represented by gray bars (exons) and coloured lines (introns), and the number above each intron is its intron phase. Three highly conserved introns in plant *PRX* genes as identified by Mathé et al. (2010) are shown in red, blue and pink, respectively. *PpPRX* genes are grouped according to clade in the Maximum-Likelihood phylogenetic tree (Fig. 2). Architecture of *PRX* genes from vascular plants and algae are given for comparison. PNP, peanut *PRX1*; HRPC, horseradish isozyme C1; KnitPrx, *Klebsormidium nitens PRX*; SsPrx03, *Spirogyra* sp. *PRX03*; CatmPrx, *Chlorokybus atmophyticus PRX*; CbraPrx01, *Chara braunii PRX01*.

In general, *PpPRX* genes in the same phylogenetic clade share identical gene architecture. For examples, *PpPRX* genes in clade C6 share the conserved ‘001’ introns except for *PpPRX47* that lost intron 3. All four *PpPRX* genes in clade F1 lost intron 3 (Fig. 1). Gene architecture was identical in each pair of sister genes at the tip of the tree (e.g., *PpPRX34*−*PpPRX39*, *PpPRX40*−*PpPRX42*, etc.). One notable exception was the *PpPRX11*−*PpPRX19* pair in clade C4, as PpPRX11 gained an intron in exon 3.

### 3.4. Chromosomal locations and gene duplications

In Fig. 3, the evolutionary history of the 27 extant chromosomes is illustrated using colours and shades, as in Lang et al. (2018). Forty-nine *PpPRX* genes and two pseudogenes were distributed unevenly among the chromosomes. Chromosomes 3 and 26 harboured the most genes (8 and 6 respectively), while seven chromosomes had none. Seven descendants of one particular anc7 chromosome (green shades in Fig. 3) were disproportionally populated as a total of 30 genes were located on them, many of them in tandem arrays.

**Fig. 3.**
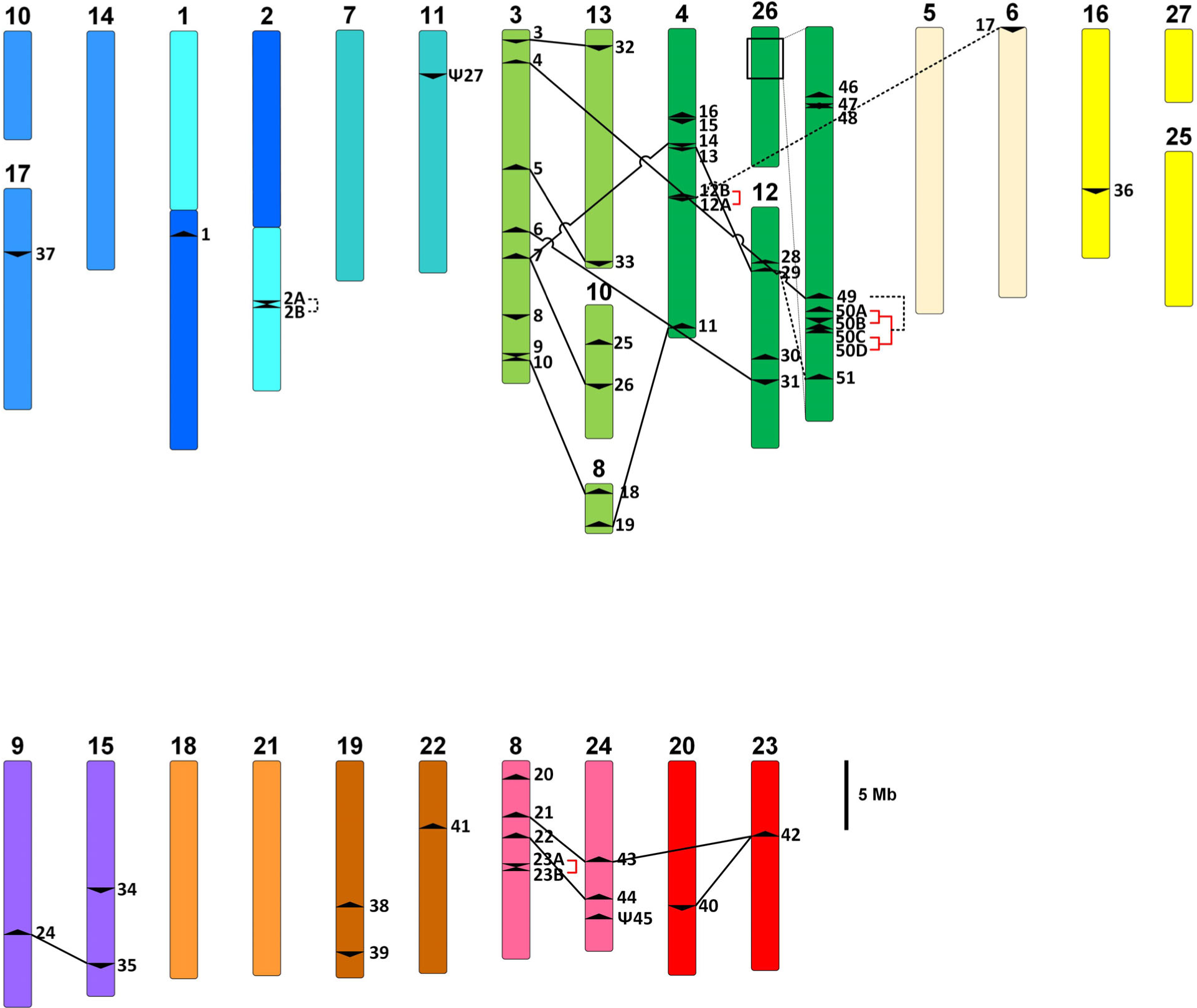
Chromosomal locations of *P. patens PRX* genes and proposed duplication pairs. A scale diagram of the *P. patens* genome was adapted from Fig 5c in Lang et al. (2018). Colours and shades of chromosomes reflect the evolution of the genome from seven ancestor (anc7) chromosomes through two WGDs and chromosomal fission or fusion events. For example, the extant chromosomes 3, 13, 10, 8, 4, 26 and 12, all in different shades of green, are descendants of one of the anc7 chromosomes, while chromosomes 5, 6, 16, 27 and 25, all in different shades of yellow, are descendants of another anc7 chromosome, and so on (for details, refer to Fig 5c in Lang et al. 2008). The location of each *PpPRX* gene was marked with arrowheads depicting forward or reverse orientation of the gene. Tandem duplications were depicted by red square brackets, segmental duplications by broken black lines, and WGDs by solid black lines. Sections of chr8 and chr26 were zoomed in for clarity.

A total of 22 duplication pairs were further analyzed (Table 2). Four genes in clade B of the ML phylogenetic tree (Fig. 1) were descendants of a single ancestral gene that was present on the red anc7 chromosome. A plausible scenario was as follows. An ancestral gene was duplicated in WGD1 to give *PpPRX42*−*PpPRX43* pair. Each gene was once again duplicated in WGD2 to give the *PpPRX40*−*PpPRX42* and *PpPRX21*−*PpPRX43* pairs. Two companion gene pairs duplicated during WGD1 were found in the immediate vicinity (<50 kb) of the *PpPRX42*−*PpPRX43* pair. They were a pair of putative *STRUCTURAL MAINTENANCE OF CHROMOSOMES* genes (Pp3c23_8100 and Pp3c24_10590) and a pair of genes encoding putative glyceraldehyde-3-phosphate dehydrogenases (Pp3c23_8160 and Pp3c24_10620) (Table 2). Similarly, at least one companion gene pair was found for 4 other WGD pairs. *PpPRX2A* and *PpPRX2B*, the two copies of *PpPRX2* with identical nucleotide sequences, also had a companion gene pair and were therefore considered as segmental duplicates. Interestingly, the companion pair, Pp3c2_28870 and Pp3c2_29010, had only 61% nucleotide sequence identity. Other gene copies were products of tandem duplications (TD).

**Table 2.**
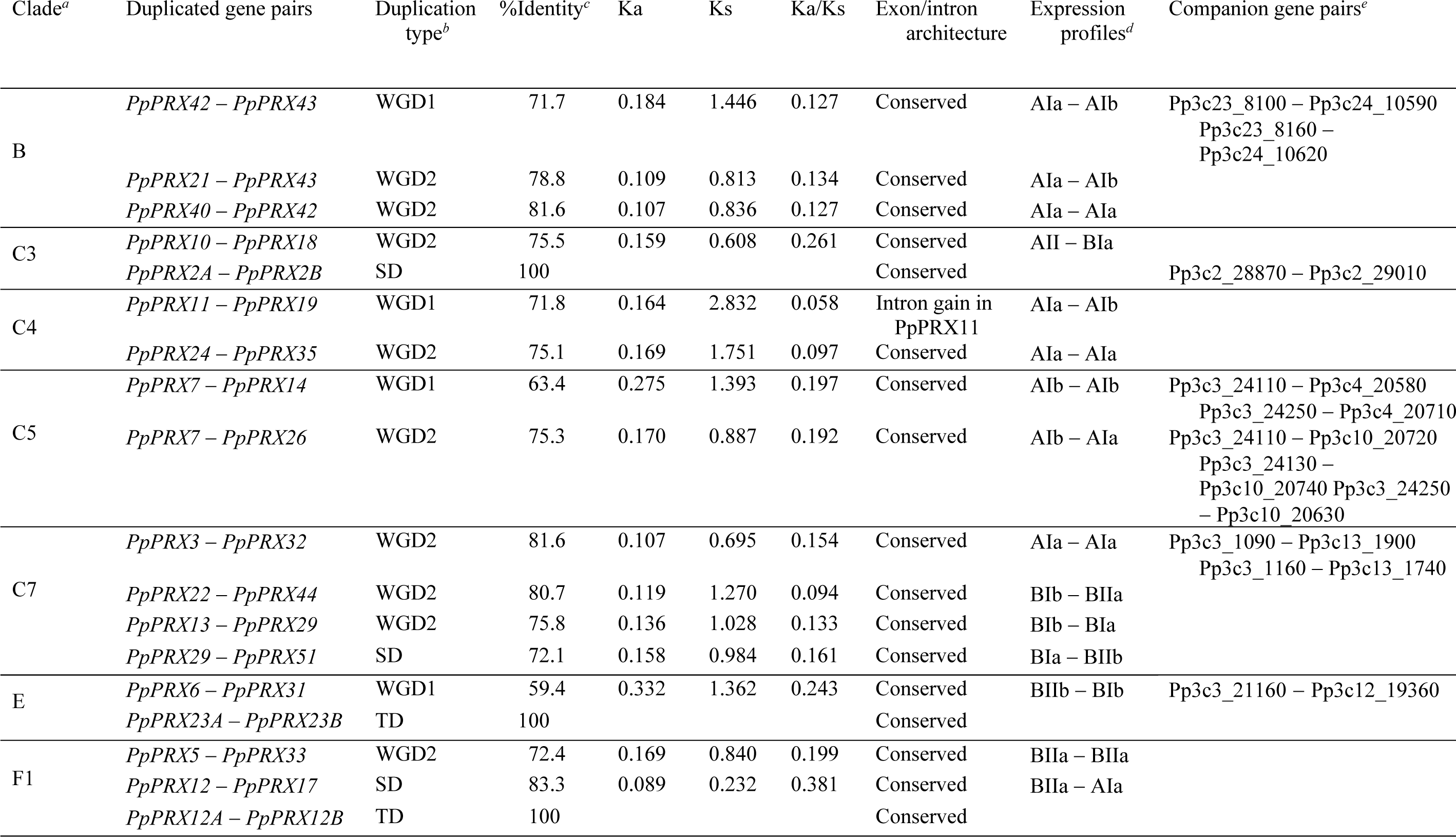
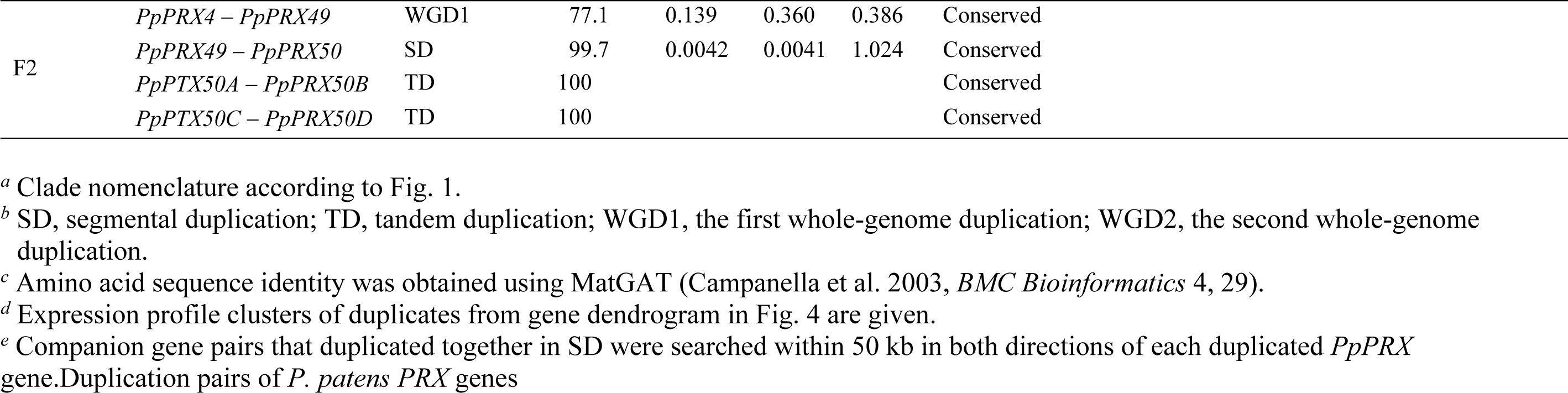
Duplication pairs of *P. patens PRX* genes. Duplication pairs from *P. patens* PRX genes are listed according to their phylogenetic clade. Their duplication type, amino acid sequence identity, Ka/Ks ratio, exon-intron architecture conservation and companion gene pairs are listed. Their expression profiles are also compared.

All duplication pairs have been under purifying selection (Ka/Ks < 1), except *PpPRX49*−*PpPRX50*, whose Ka/Ks value of 1.024 suggested neutral selection (Table 2).

Although not included in our detailed analysis due to their ≤70% sequence identity, 7 more gene pairs were considered to be possible duplicated pairs. The gene pairs, their clade name (Fig. 1), sequence identity (%) and mode of duplication are as follow. *PpPRX1*−*PpPRX38* (clade C2, 62.8%, segmental duplication (SD)), *PpPRX4*−*PpPRX25* (clade F2, 64.9%, WGD2), *PpPRX9*−*PpPRX15* (clade C6, 57.1%, WGD1), *PpPRX16*−*PpPRX48* (clade C6, 62.7%, WGD2), *PpPRX20*−*PpPRX21* (clade B, 67.5%, SD), *PpPRX20*−*PpPRX34* (clades B−A, 61.0%, SD), *PpPRX34*−*PpPRX39* (clade A, 59.1%, SD).

### 3.5. Gene expression profile

Expression data of *PpPRX* genes from different moss tissues reported by Ortiz-Ramírez et al. (2016) are presented in Fig. 4. Expression data from comparable tissues from other datasets accessible in PEATMoss were in general agreement presented (data not shown).

**Fig. 4.**
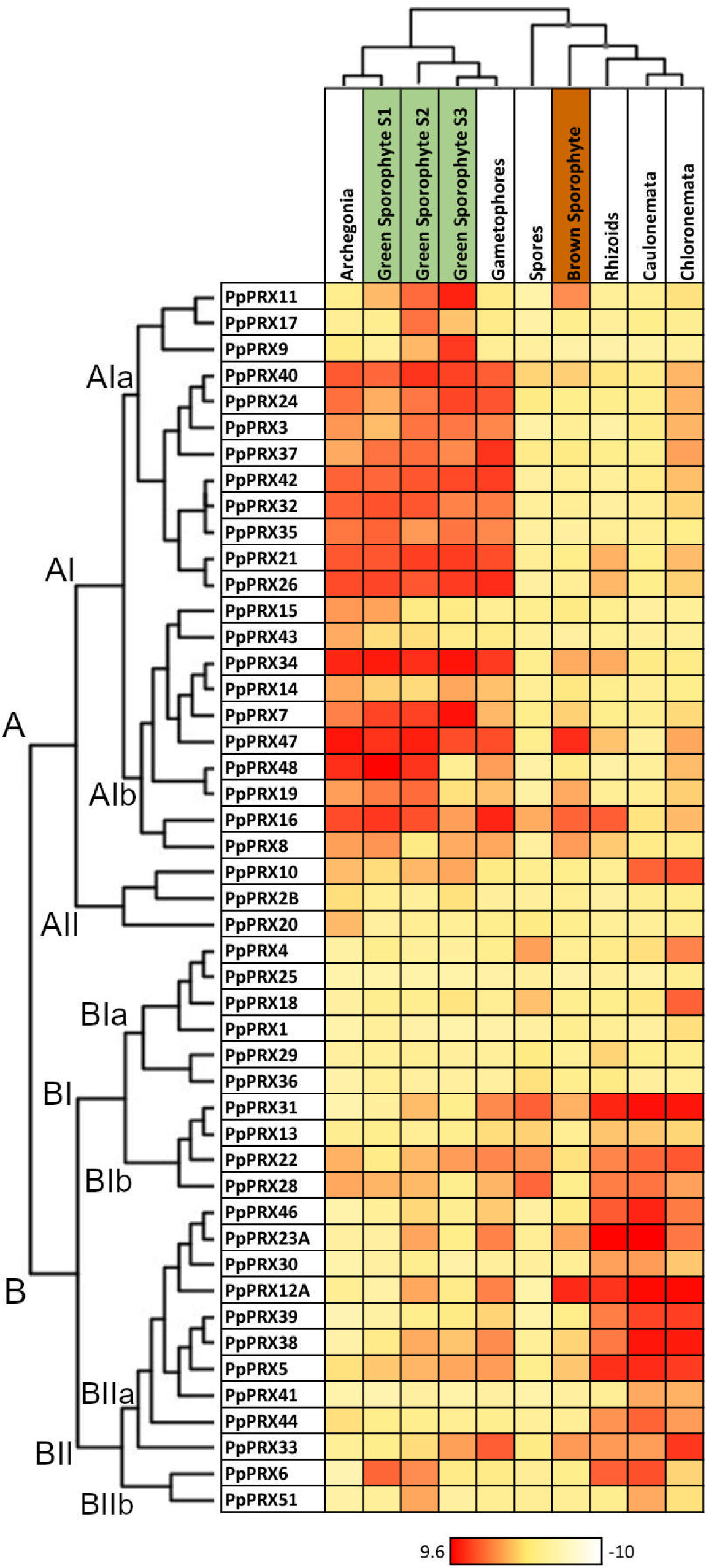
Heatmap of *P*. *patens PRX* gene expression. Expression values of *P. patens PRX* genes in each reproductive stage is represented by the colour gradient from white to red. Dendrograms show groupings of genes or tissues with similar expression patterns. The heatmap was created from log2-transformed NimbleGen microarray data from Ortiz-Ramirez et al. (2016).

As shown in the dendrogram above the heatmap in Fig. 4, expression profiles of *PpPRX* genes in different tissues were grouped into two clusters. One cluster comprised archegonia, developing green (S1, S2, S3) sporophytes and gametophores, and the other cluster comprised chloronemata, caulonemata, rhizoids, mature brown sporophytes, and spores. *PpPRX* genes were also grouped into two gene clusters according to their expression profiles, as shown in the gene dendrogram on the left side of the heatmap. Most genes in cluster A were relatively highly expressed in archegonia, green sporophytes and gametophores, while most genes in cluster B were relatively highly expressed in rhizoids, protonemata or spores. Expression levels are reported here as high (> 3.1), moderate (−3.6 to 3.1) or low ≤ −3.6), the top, middle and bottom tertiles, respectively.

Different *PpPRX* genes exhibited distinct tissue expression profiles. No gene exhibited either high or low expression in all tissues. A few genes were highly expressed in a relatively wide range of tissues. *PpPRX47* exhibited high expression in all tissues except spores and caulonemata; *PpPRX16* except in caulonemata; *PpPRX22* except in S1 green and mature brown sporophytes; *PpPRX28* except in S3 green and mature brown sporophytes; PpPRX5 except in archegonia and spores. Certain genes were highly expressed in only a few particular tissues. *PpPRX9* and *PpPRX17* were expressed highly only in S2 and S3 green sporophytes; *PpPRX15* and *PpPRX20* in archegonia and S1 green sporophytes; *PpPRX43* in S1 green sporophytes; *PpPRX4* and *PpPRX18* in spores and chloronemata; *PpPRX41* in protonemata; *PpPRX44* in rhizoids and protonemata; *PpPRX51* in S2 green sporophytes and caulonemata.

Duplicated gene pairs generally exhibited similar expression profiles. For example, *PpPRX40*−*PpPRX42*, *PpPRX24*−*PpPRX35* and *PpPRX3*−*PpPRX32*, all WGD2 duplicated pairs, belonged to gene cluster AIa and were expressed highly in archegonia, sporophytes and gametophores. There were exceptions. While *PpPRX10* of gene cluster AII was highly expressed in sporophytes and protonemata, its duplicate PpPRX18 belonged to cluster BIa and was expressed highly in spores and chloronemata. Also, PpPRX12 and its segmental duplicate PpPRX17 showed distinct expression profiles (Table 2).

### 3.6. Phylogenetic analysis of embryophyte PRX sequences

Diverse *in planta* functions reported for 62 embryophyte PRXs were grouped into five broad categories, including lignin metabolism and ROS production/oxidative burst (Table S1). An ML tree of 247 PRXs comprising (1) 62 PRXs with reported *in planta* functions, (2) phylogenetically distinct PRXs from eight plant species representing major embryophyte lineages, and (3) all PpPRXs is shown in Fig. 5. We constructed a few more ML trees using different sequence sets and tree models, and consistently made the following observations.

**Fig. 5.**
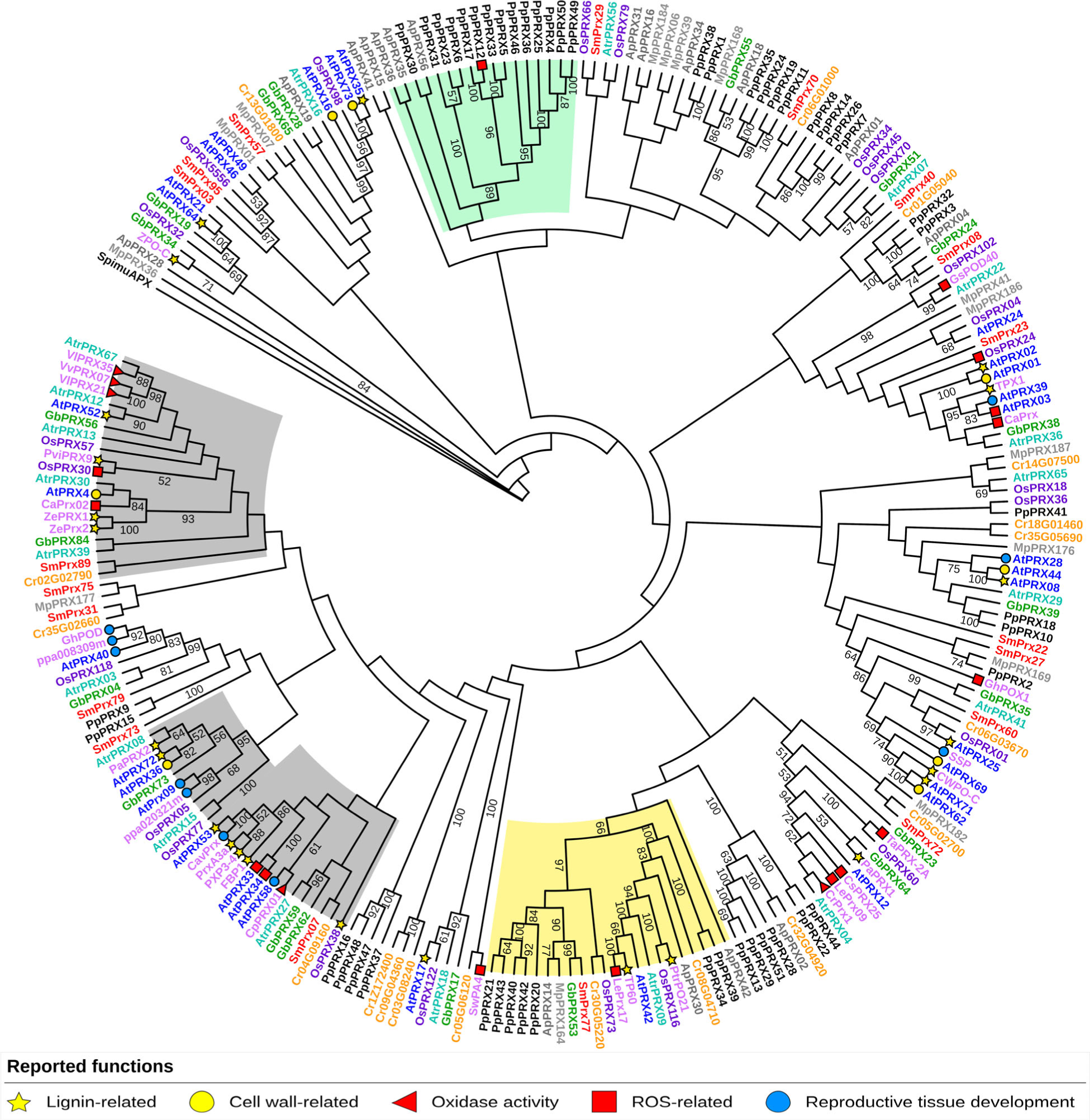
Maximum Likelihood phylogenetic tree of selected embryophyte peroxidases. The tree consisting of select PRX sequences from *Anthocerus punctatus* (ApPRX, dark grey)*, Marchantia polymorpha* (MpPRX, light grey)*, Selaginella moellendorffii* (SmPRX, red)*, Ceratopteris richardii* (orange)*, Ginkgo biloba* (GbPRX, green)*, Amborella trichopoda* (AtrPRX, teal)*, Oryza sativa* (OsPRX, purple)*, Arabidopsis thaliana* (AtPRX, blue)*, P. patens* (PpPRX, black), and PRXs with reported functions (pink) was constructed using the WAG+G+I+F substitution model for amino acids and rooted with an algal ascorbate peroxidase from *Spirogloea musicola,* SpimuAPX. Values at each node represent bootstrap support above 50% from 1000 bootstrap replicates. The clade highlighted in yellow represents an embryophyte-spanning clade. *In planta* functions, grouped in five common categories (Table S1), are indicated with different symbols. Gene IDs and sequences of PRXs from the eight plant species in the tree are provided in Table S2.

First, little clustering of the reported PRX functions was observed. PRXs with lignin metabolizing activity were found scattered in most major clades. Likewise, PRXs with ROS-related activities were present in multiple clades. Local clustering of putative orthologues or paralogues was also observed. AtPRX40, GhPOD and ppa008309m, all implicated in male reproductive processes, were found in a tip clade together, while LePrx09, CsPRX25 and TaPRX-2A, all implicated in stress response, were also found closely clustered (Fig. 5).

Second, one clade was found to contain PRX sequences from *P. patens* and each of the other eight lineage-representing species included in the tree reconstruction. This land plant-wide clade (yellow-shaded in Fig. 5) contained seven PpPRXs. PpPRX34 and PpPRX39, which were in the early divergent clade A in the PpPRX ML tree were also found in this ‘land plant-wide clade,’ along with the five PpPRXs in clade B (Fig. 1). Also found in the clade were three angiosperm PRXs with reported functions: poplar PrtPO21 and tobacco TP60, both with lignin polymerization activity, and tomato LePrx17 involved in pathogen defense (Table S1 and references therein). OsPPX73 and OsPRX116, and AtPRX42 had previously been reported as candidate ancient PRXs in rice and *Arabidopsis*, respectively (Tognolli et al. 2002; Passardi et al. 2004) (see Discussion).

Third, there were taxa-specific clades: a bryophyte-specific clade and two tracheophyte-specific clades (shaded in green and gray, respectively) were found in the tree. Other clades contained PRXs from most plant species, but lacked PRXs from a few species, suggesting lineage-specific gene losses.

## 4. Discussion

### 4.1. With conserved characteristics and relatively fewer genes, the *PpPRX* family is a good model for further study

Amino acid residues characteristic of PRX proteins at the heme-binding site and two Ca^2+^-binding sites were highly conserved in all 49 PpPRXs. A total of 9 PpPRXs are predicted to lack one of the four conserved disulfide bonds. This is not unusual. However, the frequency of mutation (9 of 49, 18%) was higher than <5% reported by Mbadinga Mbadinga et al. (2020) who counted 41 PRXs missing at least one Cys residue out of 959 PRXs analyzed. The disulfide bond Cys6−Cys7 is close to the substrate binding channel and may affect substrate selectivity (Mathé et al. 2010). PpPRX36, which is a singleton in clade F2 in the ML tree of the PpPRX family (Fig. 1) and is expressed moderately across the tissues (Fig. 4), is the only one with a mutation of Cys6.

Gene architecture of four exons and three ‘001’ introns has been widely conserved in embryophyte *PRX* genes and it was suggested that this intron pattern was established in the last common ancestor of embryophytes (Yan et al. 2019). Our finding of the same intron patterin some streptophyte algal *PRX* genes (Fig. 2) suggested that this particular *PRX* gene architecture was already present in the common ancestor of the streptophyte algal and embryophyte lineages. Intron loss was more frequent than intron gain during plant evolution (Roy and Penny 2007). Similarly, there were more intron losses than gains in *PpPRX* genes.

In this study, we examined all genes annotated as class III PpPRXs in the databases to select 49 *PRX* genes, including two *PRX* genes that have not been reported previously. Due to the stringent criteria we employed, the total number is less than what has been reported in the literature (e.g., 57 *PpPRX* genes in Mbadinga Mbadinga et al. 2020; 60 in Yan et al. 2019). In addition to the two pseudogenes (Table 1), we found 5 gene fragments that would produce short proteins of 86 to 177 amino acid residues (data not shown). Due to massive and recent duplications, *PRX* families in vascular plants are larger; there are 73 *PRX* genes in *Arabidopsis*, 93 in poplar, 102 in potato, and 119 in maize. The large number of genes and high functional redundancy in vascular plants make it challenging to study the functions of tracheophyte *PRX* genes (Cosio and Dunand 2009). With its smaller size, the *PpPRX* family may serve as a model system for functional studies of plant *PRX* genes.

### 4.2. The *PpPRX* family expanded mainly through whole-genome duplications

Among the 18 non-tandem duplications observed in the *PpPRX* family, 14 were attributed to WGDs and 4 to SD (Table 2). There were only 4 TDs, accounting for 18% (4 of 22) of total duplication events. This was in sharp contrast to the larger contribution of TD in the expansion of vascular plant *PRX* families. There were 57 SD and 26 TD events in potato, and 16 SD and 12 TD events in maize. In the model C4 grass *Setaria viridis*, 70% of *PRX* genes, 108 of 154 genes, were produced through TD events (Simões et al. 2023). It was argued that genes encoding proteins that are more loosely connected in a network, such as *PRX* and others involved in stress responses in plants, are less restricted by gene dosage effects and hence would be retained more frequently after TD events. For example, the poplar vacuole *PRX* genes expanded by TD (Ren et al. 2014, and references therein). The biological significance of the relative rarity of TD events in the *PpPRX* family is unclear.

Duplication mode affects divergence of expression profile. In *Arabidopsis*, poplar and pear, genes produced through WGD were found to exhibit a lower divergence of expression profile than other duplicated genes (Casneuf et al. 2006; Rodgers-Melnick et al. 2012; Qiao et al. 2018). This may explain why most WGD-produced *PpPRX* genes showed similar expression profiles.

Purifying selection has been a common feature in the evolution of plant *PRX* genes. Duplicated *PRX* genes in maize (Wang et al. 2015), pear (Cao et al. 2016), wheat (Yan et al. 2019), *Brachypodium distachyon* (Zhu et al. 2019), grapevine (Xiao et al. 2020), carrot (Meng et al. 2021), birch (Cai et al. 2021), tobacco (Cheng et al. 2022), and cotton (Chen et al. 2022) were all predominantly subjected to purifying selection, indicating that the gene family has undergone relatively conservative evolution with stable structure and function. The *PpPRX* family was not an exception and all duplicated pairs but one were subject to negative selection.

### 4.3. Searching for ancient and conserved *PRX* functions

*PRX* genes were first thought to have evolved in the lineage leading to land plants (embryophytes) and researchers have long sought to learn how ancestral *PRX* genes may have facilitated terrestrialization of ancient plants. It was speculated that ancient *PRX* genes might have functioned to mitigate UV-induced oxidative stresses or to contribute to formation of novel cell wall structures (Passardi et al. 2004).

The single embryophyte-wide clade in our phylogenetic tree (Fig. 5) included two moss sequences, PRX34 and PRX39. OsPRX73 and OsPRX116 had already been shown to be phylogenetically closer to a liverwort PRX than to any other rice PRX (Passardi et al. 2004). Likewise, the sequence of AtPRX42 was found to be ∼80% identical to those of PRXs from cotton, soybean and tobacco and 57% identical to its closest paralog, AtPRX21 (Kjaersgård et al. 1997; Tognolli et al. 2002). AtPRX42 and AtPRX21 were suggested to have conserved the sequence and function of an ancestral embryophyte PRX gene (Kjaersgård et al. 1997; Tognolli et al. 2002). We found that PpPRX34 and PpPRX 39 were 42% and 36% identical, respectively, to AtPRX42. That is, AtPRX42 showed higher % identity to its *P. patens* orthologues than to any other AtPRX except AtPRX21.

*AtPRX42* (At4g21960) and *AtPRX21* (At2g37130) are constitutively expressed in all tissues examined, including 2-day-old germinating seeds (Kjærsgård et al. 1997; Valério et al. 2004). A recent study showed that AtPRX42, one of the most predominant AtPRXs in the *Arabidopsis* stem, was localized in nonlignifying xylem parenchyma and phloem cells and proposed to be involved in various stress responses (Hoffmann et al. 2020). This agreed remarkably well with the functions proposed by Passardi et al. (2004) for ancient PRXs: protection of primary cell walls from oxidative stresses. We were not able to find any functional study of OsPRX73 (LOC_Os05g14260) and OsPRX116 (LOC_Os07g49360). Both genes were annotated as ‘Stress Response’ (GO:0006950) genes in the Rice Genome Annotation Project (http://rice.uga.edu/). *OsPRX73* is expressed at low levels in anther, leaf, panicle, root, seed, and shoot, while *OsPRX116* is expressed at a high level in panicle and moderately in leaf, root, seed and shoot (http://expression.ic4r.org/). Interestingly, the two *PpPRX* genes in the clade showed expression profiles that are complementary, suggesting subfunctionalization of the genes after duplication. *PpPRX34* of expression cluster AIb (Fig. 4) was highly expressed in archegonia, sporophytes and gametophores, whereas *PpPRX39* of BIIa was mainly expressed in rhizoids and protonemata.

The land plant-wide clade in the ML tree (Fig. 5) is by no means complete. Further studies with more sequences from diverse taxa and their functional characterization will be required. Current data suggested that stress response was the main function of ancient and conserved PRXs in the land plant-wide clade and gene duplications and subfunctionalization (OsPRX73 and OsPRX116) or neofunctionalization (TP60 and PrtPO21 in lignification) subsequently occurred in different lineages. In *P. patens*, at least six duplications occurred to create the seven PpPRXs in the clade. Functional studies on some of these PRXs should be informative.

### 4.4. Candidate *PpPRX* genes for future studies

PRX can oxidize a variety of substrates or generate ROS *in planta*, and exhibits substrate promiscuity *in vitro* (Passardi et al. 2005). In addition, functional redundancy and compensation among *PRX* genes have made it difficult to study the biological functions of individual *PRX* genes. It has also been difficult to predict PRX functions based on sequence similarity and phylogeny (Cosio and Dunand, 2009; Mathé et al, 2010). This is partly due to functional divergence of duplicated genes as in the case of *AtPRX72* in lignification and *AtPRX36* in cell wall loosening and seed germination (Fig. 5; Table S1). Further, some PRX functions (*e*.*g*., lignin polymerization) appear to have evolved multiple times through neofunctionalization in a gene family. This was observed in Arabidopsis, poplar and *Zinnia elegans* (Fig. 5; Table S1).

Nonetheless, recent studies have pointed out that sets of amino acid residues, mostly at or close to the enzyme active site, have been positively selected for and thus might be useful in function prediction (Ren et al. 2014; Kupriyanova et al. 2015; New et al. 2023). The present study also showed phylogenetic clustering of *PRX* genes from different taxa, and this could be used to predict functional similarity among the genes. As examples, GhPOD (Malvaceae) and ppa008309m (Rosaceae), which were implicated in male reproduction, belonged to the same clade as AtPRX40 (Brassicaceae) (Fig. 5). AtPRX40 functions as an extensin peroxidase during anther development (Jacobowitz et al. 2019), and GhPOD, ppa008309m and some other genes in the same clade may share the same function. No bryophyte *PRX* gene has been shown to be involved in sporophyte development, although ROS was required in spore wall formation (Rabbi et al. 2020). The three *PpPRX* genes in the AtPRX40 clade (*PpPRX9, PpPRX15, PpPRX37*) are highly expressed in developing sporophytes (Fig. 4), and *PpPRX9*, in particular, was up-regulated in the sporophyte transcriptome and coexpressed with sporopollenin biosynthetic genes (O’Donoghue et al., 2013). Functional characterization of these *PpPRX* genes will reveal the extent of functional similarity in the AtPRX40 clade. Similarly, studies on *PpPRX34* and *PpPRX39* in the ancient land plant-wide clade will be informative regarding ancient functions of embryophyte *PRX* genes and the extent of gene functional similarity in the land plant-wide clade.

In conclusion, 49 *PpPRX* genes were identified and their sequences and gene architecture were analyzed. WGDs were mostly responsible for the family expansion, and duplicated genes were under purifying selection, while generally sharing similar expression profiles. A phylogenetic reconstruction with PRX sequences from the major lineages of land plants revealed an ancient land plant-wide clade. Due to the relatively smaller size of the gene family and simpler morphology and physiology of the moss, the *PpPRX* gene family may prove to be a fertile system for functional studies of PRX genes. *PpPRX34* and *PpPRX39* are proposed as candidate genes to test this premise.

## Author contributions

Conceptualization, funding acquisition: D-YS

Data curation, formal analysis, investigation, methodology, validation: VA, FR, TM, WB, EB, D-YS

Writing – original draft: VA, EB, D-YS Writing – review & editing: EB, D-YS

## Data availability

All data generated or analyzed during this study are included in this published article (and its supplementary information files).

