## Supplementary material for "Genome-wide analysis of the class III peroxidase gene family in *Physcomitrium patens* and a search for clues to ancient class III peroxidase functions": Fig. S1

**Table S1** Class III peroxidases with reported *in planta* functions
**Table S2** Embryophyte PRXs used to reconstruct the Maximun Likelihood phylogenetic tree: Gene ID and sequence
(Supplementary tables are presented in separate Excel worksheets.)

**Fig. S1** Amino acid sequence alignment of class III peroxidases from *P. patens*

**Fig. S1 Amino acid sequence alignment of class III peroxidases from *P. patens***

Residues at the heme binding site are highlighted in pink with proximal (*P*) and distal (*D*) residues indicated on the tops of the columns, and $Ca^{2+}$ binding residues (side chains only) are highlighted in yellow. The eight Cys residues (in cyan) that form four disulfide bonds are numbered 1 to 8 on the tops of the columns. Disulfide forming pairs are Cys1 and Cys4, Cys2 and Cys3, Cys5 and Cys8, and Cys6 and Cys7. Exon-intron junctions are highlighted in green. Class III PRX sequences from four algae and two vascular plants are also aligned for comparison. CbraPrx01 from *Chara braunii* S276 (Charophyceae), CatmPrx from *Chlorokybus atmophyticus* CCAC 0220 (Chlorokybophyceae), SsPrx03 from *Spirogyra* sp. (Zygnemataceae), KnitPrx from *Klebsormidium nitens* (Klebsormidiophyceae), HRPC from horseradish (isozyme C1, P00433.2), and PNP from peanut (cationic peroxidase 1, NP_001363143). Algal PRX sequences were obtained from RedOxiBase (https://peroxibase.toulouse.inra.fr/) and PhycoCosm (https://phycocosm.jgi.doe.gov/phycocosm/home).

```
PpPRX1      ---------MG------GGPTAM------------LSVLALLVASL--------------    19
PpPRX38     ---------MR-TTNMAAALTVR------------LPLLCILLMCL--------------    23
PpPRX4      ---------MG-KWSSKAL----------------ATIFVTLAIAM--------------    20
PpPRX49     ---------MG-RWPSPAL----------------LAIFVTIALAM--------------    20
PpPRX50     ---------MG-RWPSPAL----------------LAIFVTIALAM--------------    20
PpPRX25     ---------MG-RWASTGF----------------LAIFITIAIAI--------------    20
PpPRX46     ---------MA-SAWYVR-----------------LCAVV-IILSLD--------SFVSA    24
PpPRX6      ---------MD-HPHRA------------------SLLALLIVVVL--------------    18
PpPRX23     ---------MA-HHAM-------------------RATVIAVVLVA--------------    17
PpPRX31     ---------MA-RILRAS-----------------CIFALVCVIAI--------------    19
PpPRX36     ---------MS-------------------------------------------------     2
PpPRX5      ---------MS-IYQGGL-----------------VAALLAVAISL--------------    19
PpPRX33     MSEVNPSKRMP-NHRRGL-----------------ATALLTIAFVF--------------    28
PpPRX12     ---------MA-AFDPMGASKL-------------QVLLMTVAVAM--------------    23
PpPRX17     ---------MG-AFDPMGPSKM-------------QVLLMTVALAM--------------    23
PpPRX7      ---------ME-MRLHSKK----RKPAL------WTVMAMMIAQLI--------------    26
PpPRX26     ---------ME-SRHNLLQ----RRPTL------WTAVVLVILQLA--------------    26
PpPRX14     ---------ME-RRCKHLH----WKAFLVT-----TPALLWFAMLL--------------    27
PpPRX8      ---------ME-SAAGRT-----------------QRALAVWLVIA--------------    19
PpPRX11     ---------MA-ISCR-------------------VWRLLLALCTL--------------    17
PpPRX19     ---------MA-GYR--------------------IWQLLFAVWAV--------------    16
PpPRX24     ---------MG-GLDFSRRSSAAE-----------SCFVVVIALSL--------------    25
PpPRX35     ---------MA-GLRYGRTNRAT------------VAMASLLLAAL--------------    24
PpPRX9      ---------MA-IYKSHSPQCA-------------VWMALTLVTCV--------------    23
PpPRX15     ---------MH-SGDRMRCLTRC------------SIFKAVAAMML--------------    24
PpPRX37     ---------MD-ACYSSST----------------VVLLLLVLSCM--------------    20
PpPRX16     ---------MK-AMGASGAVM--------------AVRVLSVVVLL--------------    22
PpPRX48     ---------MA-ISRAR-------------------ARVVVALILV--------------    17
PpPRX47     ---------MA-SFR---------------------AGAAVSLCLM--------------    15
PpPRX2      ---------MS-RSTKIGSDSQAPA----------PWLLTILIASS--------------    26
PpPRX10     ---------MA-EQRLRGG----------------AWGVVLLLVRL--------------    20
PpPRX18     ---------MA-AQRGRGG----------------AFHLLLLLVSL--------------    20
PpPRX30     ---------MA-RIHRDVSVLGSM-----------ALVALCLYCGV--------------    25
PpPRX3      ---------MA-HFQIGEGFKWEAAF---------FCCWVLCWVAV--------------    27
PpPRX32     ---------MA-HFQIQERFIWGAVL---------CCCFVLCLFSA--------------    27
PpPRX13     ---------MK-SFLLIGV----------------IYGLCALSASAWECSFNGRRKVLQV    34
PpPRX29     ---------MKSSLLLIGV----------------ICSLLALSASAEECPFSGRRNLLQA    35
PpPRX51     ---------MD-------------------------------------ATFFKLEHLIQP    14
PpPRX28     ---------MK-NLGVIGVVYG-------------LMALATLQVASAWCPFAGRRSLLQA    37
PpPRX22     ---------MA-RIRFQQT----------------LIVLLSLAAFF--------------    20
PpPRX44     ---------MS----ILQT----------------LALLLCLAVSF--------------    17
PpPRX20     ---------ME-------------------------WGSVLILLLL--------------    12
PpPRX21     ---------MG------------------------RLLALELLTAA--------------    13
PpPRX43     M--------MG-IWTAF------------------KTLAAALLLVA--------------    19
PpPRX40     ---------MA-SWTVT------------------LLVGAAVAVVV--------------    18
PpPRX42     ---------MA-GWTAA-----------------LLGAAAAGVYMV--------------    19
PpPRX34     ---------MD-RL---------------------AALLIALFCLL--------------    15
PpPRX39     ---------MA-RLGS-------------------FVAVLVLVCAA--------------    17
PpPRX41     ---------MR-RDKHLRL----------------LVTYIIFLLAAN--SFS---SLLGE    29

CbraPrx01   ---------MN-RRVTWRSGLRKALLLQLLLTVWWLICLSDLCVVA-----------AKE    39
CatmPrx     ---------MA------------------------SAVVVLLLLLS--------------    13
SsPrx03     ---------MG-SESKPSY----------------FLALRTLDTQH--------------    20
KnitPrx     ---------MA-PPTF-------------------SWHILLTLLAV--------------    17
HRPC        ---------MHFSSSSTLF----------------TCITLIPLVCL--------------    21
PNP         ---------MA------------------------LPISKVDFLIF--------------    13
```

```
PpPRX1     -----------------------------------------AALSTTVQAQ-LVENFY--    34
PpPRX38    ------------------------------------------ASVTTIQAQ-LSTNFY--    39
PpPRX4     ----------------------------------------NSITPAVAHSG-LKVGFY--    37
PpPRX49    ----------------------------------------NSIIPAAAHTG-LKVGFY--    37
PpPRX50    ----------------------------------------NSITPAAAHTG-LKVGFY--    37
PpPRX25    ------------------------------------AIVMNLIIPAGAHKG-LEVGFY--    41
PpPRX46    SKYD--------------------------------KVHHNSGTHHVPDGG-LRDNYY--    49
PpPRX6     -----------------------------------------AVSVSSAEGQ-LVYRYY--    34
PpPRX23    -----------------------------------------MLAVTGVDAT-LRYGFY--    33
PpPRX31    -----------------------------------------SLSVNQVDA--LDYNYY--    34
PpPRX36    ----------------------------------------TIPVNTEDGAD-LHYDFF--    19
PpPRX5     ----------------------------------------TCLSSHAEAQG-LAVGYY--    36
PpPRX33    ----------------------------------------VCLSSQAEAQG-LTVGFY--    45
PpPRX12    --------------------------------------FFLSGEVEANSYN-LRPNYY--    42
PpPRX17    ---------------------------------------FLSGEVEATAYN-LRPHYY--    41
PpPRX7     -----------------------------------------FHPRMCTGGA-LRPGYY--    42
PpPRX26    -----------------------------------------LSCWMCTVDA-LQTGYY--    42
PpPRX14    -----------------------------------------VAMRLSSAEP-LRVGYY--    43
PpPRX8     -----------------------------------------QLLQIVAAQD-LQVDFY--    35
PpPRX11    ----------------------------------------GSVGMESVNAQ-FFNNFY--    34
PpPRX19    -----------------------------------------SGSGVEVSAQ-F-VGYY--    31
PpPRX24    -----------------------------------------LLVTQVRAQN-IGVGFY--    41
PpPRX35    -----------------------------------------VFQGHVQVGG-VAVGFY--    40
PpPRX9     ------------------------------------GGKILSVAATWDDSH-LTPSFY--    44
PpPRX15    -----------------------------------------VVVVKICAAE-LDVAYY--    40
PpPRX37    --------------------------------------QLVVQGSLDNQYG-LRKSYY--    39
PpPRX16    -----------------------------------AVVSTVGAEAVVASGL-LTTDYY--    44
PpPRX48    -----------------------------------------KFVILVNAQV-LTTEFY--    33
PpPRX47    -----------------------------------------TLVTMLSVDA-LTTDFY--    31
PpPRX2     -----------------------------------------IAVQHVVHGA-VQVGFY--    42
PpPRX10    ------------------------------------------MSLLHGTKA-LRVGFY--    35
PpPRX18    ------------------------------------------MCVFHGTQG-LRVGFY--    35
PpPRX30    -----------------------------------------AGAVLNYTPG-LRYDFY--    41
PpPRX3     -----------------------------------SSNAEGRAVNASTRPK-LNRLFY--    49
PpPRX32    -----------------------------------RTEAQIGVATPRIVPK-LNTLFY--    49
PpPRX13    GAPDAAPP-------QQLEAPSPSAPSSPPLPPQSTVATRPTGADVHGEPP-LDYDFY--    84
PpPRX29    GAPGAAPPA------QQGGAPAPAA-----------VGPRPVGKDVYGEPA-LDYDFY--    75
PpPRX51    RHPNRA---------LQLHRHFPHA---------------PRGKDVYGGPP-LDYDFY--    47
PpPRX28    AAP----------------APLP---------------ARPTGPDPSNWTM-LNYDFYTA    65
PpPRX22    ---------------RVGDAGCPIA---------SLGRRSLKQAPVPAQPA-LIFGFY--    53
PpPRX44    ---------------QPGDARCPLA---------ILGRRSLKQADLPPQPP-LQFGFY--    50
PpPRX20    -----------------------------------------SSVILFSEGH-LSSDYY--    28
PpPRX21    -----------------------------------------FLLAAVADAE-LRYGYY--    29
PpPRX43    -----------------------------------------MILKPVVEAQ-LQYGYY--    35
PpPRX40    -----------------------------------------TNLAAVTDAE-LQYGYY--    34
PpPRX42    -----------------------------------------LFLGTTVHAE-LQYGYY--    35
PpPRX34    -----------------------------------------ATVLKVESEG-LVYDYY--    31
PpPRX39    -----------------------------------------SFVNVADSAG-LVNNFY--    33
PpPRX41    AASE------------KNT-----KKGG-----DQNGDQNVAAVPPAQASK-LRYNHY--    64

CbraPrx01  EESEVVHHVITISGKRRASGRLPFSSSS-----SSSSSSKPKYASTAVKGK-IDYSLYLP    93
CatmPrx    -----------------------------------------SFASQAAAAS-LTYKHYLK    31
SsPrx03    ------H----------------------------------SKILRNRDTR-LWKNHFVQ    39
KnitPrx    ------------------------------------SWLNQTVIAISIVSP-LNVTYY--    38
HRPC       -----------------------------------------ILHASLSDAQ-LTPTFY--    37
PNP        -----------------------------------------MCLIGLGSAQ-LSSNFY--    29
```

```
                        1                                    D   DD 2      3
PpPRX1      ---R-TSCP--SAETVITSAVNSALNRRAA---------SAAGVLRIHFHDCFV--HGCD    78
PpPRX38     ---R-STCK--DAETIISVAVTSALSRRPA---------AAAGIIRMLFHDCFV--HGCD    82
PpPRX4      ---H-HSCP--EVETIVYNSMVQSYKANHT---------VAPGVLRMAFHDCFV--RGCD    80
PpPRX49     ---R-HSCP--QVEAIVYNSMAQSTKADDT---------VAPGILRMAFHDCFV--RGCD    80
PpPRX50     ---R-HSCP--QVEAIVYNSMAQSTKADDT---------VAPGILRMAFHDCFV--RGCD    80
PpPRX25     ---R-NSCP--EVETIVYKSMAQSYKTNMT---------VAPGVLRLAFHDCFV--RGCD    84
PpPRX46     ---H-RSCP--HVEKIIFKEISKCFKADKK---------IAPGILRMSFHDCFV--RGCD    92
PpPRX6      ---K-QSCP--NVEKIIHKEVLKQFKKDPT---------IAPGILRLIFHDCFV--RGCD    77
PpPRX23     ---K-HSCP--NVESIIYKAMKAAYEKDNT---------VAPGVLRLIFHDCFV--RGCD    76
PpPRX31     ---R-KSCP--QAESIIFREVQRYFKKDPT---------VAPGLLRLIFHDCFV--RGCD    77
PpPRX36     ---E-HNCP--EVENIVCNPIYESYLKNST---------IAPGVLRMAYHDCFV--RGCD    62
PpPRX5      ---T-NSCP--TVETIIYNSMWDSYTRDPT---------TAPGVLRLAFHDCFV--RGCD    79
PpPRX33     ---E-NSCP--TVEAIIWESMRNSYNQDPT---------VAPGVLRLSFHDCFV--RGCD    88
PpPRX12     ---S-GKCGKYDVESIIYNEIAKAFQQDNG---------VAPGLVRMAFHDCFV--RGCD    87
PpPRX17     ---R-GKCGRHDVEKIIYNAVAQAFRQDNG---------VAPGIVRLAYHDYFV--RGCD    86
PpPRX7      ---A-QTCP--NAENIIRAAMEWGMQQDSG---------TAPGVLRLHFHDCFV--DGCD    85
PpPRX26     ---A-ATCP--NAEAIIRAAMERGMQEDSG---------TAPGVLRLHFHDCFV--DGCD    85
PpPRX14     ---D-LSCP--SAERIIRQAMERGMQQDQG---------IAAGVLRLHFHDCFV--EGCD    86
PpPRX8      ---G-GTCP--SAEKIVRDAVEAAVAKDHG---------NAPGLIRLHFHDCFV--RGCD    78
PpPRX11     --TT-KGCD--SAEAIVTQAVTEAFNQDPS---------VAPALIRMLFHDCFV--EGCD    78
PpPRX19     --RT-KDCG--IAEAIVTQAVTQAFNQDPS---------VAPALIRLLFHDCFV--EGCD    75
PpPRX24     ---D-QSCP--RAESIVTETVREFNSRDAT---------VPAALLRLLFHDCFV--EGCD    84
PpPRX35     ---D-QTCP--QAESIVTQTVREFNSKDPT---------TPAALLRLLFHDCFV--EGCD    83
PpPRX9      ---D-NKCP--HLQKVVSSKVEAGRRRDQR---------LPASVLRLHFHDCFV--NGCD    87
PpPRX15     ---D-FRCP--DALAIVQGGVHAAMQRDAR---------APASLLRLHFHDCFV--NGCD    83
PpPRX37     ---G-VSCP--NAEEIVTKTVTKAVKHDSR---------SAASLVRLFFHDCFV--SGCD    82
PpPRX16     ---A-RSCP--RLSSIVKEEVQKAVKVEKR---------MAASLVRLHFHDCFV--HGCD    87
PpPRX48     ---D-ESCP--EIYSIVKEEVQKAVEAEKR---------MAASLIRLHFHDCFV--NGCD    76
PpPRX47     ---A-KSCP--RIHSIVKAEIKKAVNVEKR---------MAASLIRLHFHDCFV--HGCD    74
PpPRX2      ---D-ATCS--AAESIVKGAVQSAVALDGT---------IAASIIRLHFHDCFA--QGCD    85
PpPRX10     ---N-NICP--GTETIVRQVVENRFSRDQS---------ITPALLRLFFHDCFV--TGCD    78
PpPRX18     ---T-NTCP--NAETIVTQTVQNRFRRDKT---------ITPALLRLFFHDCFV--VGCD    78
PpPRX30     ---K-NSCP--RADDIVFEQMTEIFKTKPTAQDGDFGKNVAPDLLRLHFHDCFV--RGCE    93
PpPRX3      ---S-HSCP--RLEHVVSSTMARHLQQNIA---------SGAPLLRMFFHDCAV--NGCD    92
PpPRX32     ---S-HSCP--GLQQVVTSTMARNLQQDIS---------SGAPLLRMFFHDCAV--NGCD    92
PpPRX13     ---T-KSCP--SFQQIVKTQVARAILADSL---------TPAKLLRLFFHDCFV--MGCD   127
PpPRX29     ---T-TKCP--TFLQIVKTEVAKAIAADSL---------TPAQLLRLFFHDCFV--MGCD   118
PpPRX51     ---T-TMSP--SFQQIVKSEVAKAMLADSL---------TPAQLLRLFFHDCFV--MGCD    90
PpPRX28     GLPE-NACP--SFQNIVKKEVAKATVLDSL---------TPAFLLRLFFHDCFV--MGCD   111
PpPRX22     ---D-LTCP--TLNSIIDTRMRFWVLQDIR---------TPGKVLRLFFHDCFA--AGCE    96
PpPRX44     ---D-ISCP--SLNAIIDERMRFWVLQDIR---------TPAKILRLFFHDCFA--AGCE    93
PpPRX20     ---K-STCP--NVERVVRSSLRRAFLLDPS---------APASLLRLSFHDCQV--EKCD    71
PpPRX21     ---DSLGCA--GVEDRVRTLVRRSFLTDAT---------SSAAMLRLAFHDCQVGPGGCD    75
PpPRX43     ---ETLGCR--GVENLVRTSVGLSFLTDPT---------ASAAMLRLAFHDCQVGPGGCD    81
PpPRX40     ---DRVGCF--GVENRVRTLVRRSFIADVT---------ASAAMLRLAFHDCQVGPGGCD    80
PpPRX42     ---DSLGCA--GVEDRVRTLVQRSFVADAT---------ASAAMLRLAFHDCQVGPGGCD    81
PpPRX34     ---A-NSCP--NAEKIIHDTVYKLYEKKGN---------IATSLIRYVFHDCF---DSCD    73
PpPRX39     ---R-KSCP--NAEKIIRDSIYRMYEKKGN---------IATSFIRFGFHDFF---NGAD    75
PpPRX41     ---RTTRCS--KVESVVTTEVTKRSKRDPT---------LIAALLRLFFHDCFA-NKQCD   109

CbraPrx01   --SD-HDC----LLVAIRSTAGAMFRKNAG---------LAAAALRLGFHDCMTKGGGCD   137
CatmPrx     --RQ-HYCP--QFPCIVADVIKEATRRDRI---------AAPGLLRAFFHDCFV--RGCD    75
SsPrx03     --DRMKACP--QLENLVNRAFEAQWKKDRS---------VSPAILRLFFHDALV--NGVD    84
KnitPrx     ---A-TSCP--QVHEVVRERLAVAFKEDRT---------VAGSVARLLFHDCFV--EGCD    81
HRPC        ---D-NSCP--NVSNIVRDTIVNELRSDPR---------IAASILRLHFHDCFV--NGCD    80
PNP         ---A-TKCP--NALSTIKSAVNSAVAKEAR---------MGASLLRLHFHDCFV--QGCD    72
```

```
                                 D                                    4       5
PpPRX1      ASVLIDSP-----SEKDAPPNG-SLQ--GFEVIDAAKTAIEKRCPG--IVSCADITAMAS   128
PpPRX38     ASVLIDSP-----SEKDAAPNQ-SLQ--GFDVIDEAKAAVEAKCPG--IVSCSDVLALAA   132
PpPRX4      ASVLLE----GPNTERTALFNR-GLH--GFEAVDAAKRAVESACPG--IVSAADILQFAA   131
PpPRX49     ASVLLE----GPNTERRARTNT-GLH--GFDAIDAAKRAVENACPG--VVSAADVLQFAA   131
PpPRX50     ASVLLE----GPNTERRARTNT-GLH--GFDAIDAAKRAVENACPG--VVSAADVLQFAA   131
PpPRX25     AFVLLD----GPNTERKSVLNG-GLH--GYDAVDAAKRATEKACPG--IVSAPDVLQFAA   131
PpPRX46     CSLLLK----GNNTERSSRSNA-NLH--GFEALNAAKDAVEKACPG--VVSCSDVLQYAT   135
PpPRX6      ASVLLA----GKDTERTSLTNA-NLH--GFEAIDAIKAAVEKACPN--TVSCADILAYAS   128
PpPRX23     ASVLLA----GNNTERAALNNQ-GLH--GFEAIDAVKDAVEKECPG--VVSCADILAFAS   127
PpPRX31     ASVLLS----GRRSERASAINA-RLH--GFQVIDAAKHYLEDACPR--TVSCADILAYAS   128
PpPRX36     ASLLLE----GPDSEKSHPINA-PMH--GFEAIDAAKEEVEKACPG--VVSCADVLQFAV   113
PpPRX5      ASVLLD----GVDSEKAAAVNV-NLH--GFDAIDAAKTAVEDACPG--TVSCADILQYAA   130
PpPRX33     ASVLLD----GEEAEKTAAINV-NLH--GFEAIDAAKAAVEAACPN--TVSCADILQFAA   139
PpPRX12     ASLLLD----IPNSEKTATINL-GLRASAFNAIDAAKTAVESVCPG--VVSCADVLQYAT   140
PpPRX17     ASLLLD----TPNSEKTAPINR-GLRAIAFNAIDTAKAAVESVCPG--VVSCADVLQYAT   139
PpPRX7      GSVLLE----GPTSEKTAPPNS-SLR--GFEVIDAAKAELEATCPG--VVSCADILAYCA   136
PpPRX26     GSVLLD----GPRSEKTASPNL-TLR--GYEVIDAAKADLELACSG--IVSCADILAYAA   136
PpPRX14     GSVLLD----NPNSEKTSPPNF-SLR--GFEVVDAAKADLEALCPG--VVSCADILAFGA   137
PpPRX8      ASVLLD----GPKSEKVASPNF-SLR--GFEVVDAAKAELEKQCPG--IVSCADILAFAA   129
PpPRX11     GSLLLDPTPQNPNVEKLALPNL-SVR--GYEVIDAAKMQLEKTCPR--TVSCADIVALAA   133
PpPRX19     ASILLDPSPENPNVEKRSGPNL-SVR--GYEVIDAAKTQLEKTCPL--TVSCADIVALAA   130
PpPRX24     GSLLLDPSPENPDVEKAASPNL-TVR--GYDVIDAAKARLEVECPQ--TVSCADIVALAA   139
PpPRX35     ASILLDATPQNPNIEKMAAPNL-TVR--GYEVIDGAKARLEAACPG--TVSCADIVALAA   138
PpPRX9      GSILLDDRP-GFVGEKSAAPNLNSAR--GFELIDDIKQDVEALCPD--TVSCADILTIAA   142
PpPRX15     GSNLLDDRP-GFVGEKTAAPNLNSAR--GFEIIDEIKQQLEDACPK--TVSCADIVAAAA   138
PpPRX37     GSVLLDNST-TAMSEKEARPNINTLR--GFGIIERIKESLENACSE--TVSCADILALAA   137
PpPRX16     GSILLDDTA-TFIGEKTAAANNNSAR--GFEVIDGIKAKLEEACPK--TVSCADILALAS   142
PpPRX48     GSLLLDDPILGGTGEKLSRSNLNSTR--GFEVIDTIKTRLESACPN--TVSCADLLAIAA   132
PpPRX47     GSILLDSIP-GMDSEKFAPPNDRSAR--GYEAIDAIKVALEKACPR--TVSCADILAIAY   129
PpPRX2      ASIMLT----GTGSERDAPPNL-SVR--GYGVINDAKAQLESSCPG--VVSCADIIALAA   136
PpPRX10     ASLLINSTP-TNSAEKDAGANL-TVR--GFDLIDTAKAAVERVCPG--MVSCADIIALAT   132
PpPRX18     ASLLINSTP-KNSAEKDAGANL-TVR--GYDLIDAAKAAVEKACPG--KVSCADIIALAT   132
PpPRX30     GSVLMD----KPGSEKTAPPNG-RLE--GFDAVDKIKAALEGECPG--TVSCADLLAFAA   144
PpPRX3      ASVLIDSTP-NNTAERDAIPNQ-TVR--GYHIVDDIKSQVEVMCPG--IVSCADIIALAS   146
PpPRX32     GSVLIASTP-NNTAERDAVPNL-TVR--GYDIVDDIKSQVEAMCPG--IVSCADIIALAS   146
PpPRX13     ASLLLNSTV-VNLAERDHANNF-TVD--KYSVIDSIKAELEKACQA--IVSCADTLGAAA   181
PpPRX29     ASLLLKSTK-VNLAERDHPNNF-TVD--KFTVIDAIKAELEKACPG--IVSCADILGAAA   172
PpPRX51     ASLLLKSTV-LNLAERDHPSNF-TVD--KFSVIDAIKAELEKACPG--IVSCADIVGAAA   144
PpPRX28     ASVLINSTL-LNLAEKDQTKSF-SLN--KFNVVDDIKTALEVACPG--VVSCADILAAAA   165
PpPRX22     ASILLNSTA-EFAAEKDAPISV-TLD--KFQVIEDIKSEVETACPG--IVSCADILALAA   150
PpPRX44     ASILLNSTS-TVLAEKDAPISQ-TLD--KYDVIENIKWTVETACPG--IVSCADILALAA   147
PpPRX20     ASILLDSVSNDINGERESGGNF-GIR--RLDIIDRVKQDLEKECPG--VVSCADIVAMAG   126
PpPRX21     GSIMME----GDGGEMDAGSNF-GVK--RLDIINSVKSDMEDMCPL--TVSCADIIAMAG   126
PpPRX43     GSIMVE----GNGREMDAGGNF-GVK--RLDIINSVKADLERMCPM--TVSCADIIAMVG   132
PpPRX40     GSIMIE----GNGGEMSSGNNF-GVK--RLDIINSVKADMEKMCPT--TVSCADIIAMAG   131
PpPRX42     ASIMID----EDAGEMASGNNF-GIK--RLDIINSVKADMEDNCPN--TVSCADIIAMAG   132
PpPRX34     ASVLLESSK-GVPAEKESHSQV-GMR--NGKWINNIKKAVEDSCPG--VVSCADVLALGG   127
PpPRX39     ASFFLLSAP-GKTSEKDSHSMV-GMR--NEKYVNNIKAEVEKVCPG--VVSCADILAVGS   129
PpPRX41     ASLLIKKMPKNEKTELHSGSNV-GIK--GLAFMDYLKTKVVESCNGSNIVSCADILALAA   166

CbraPrx01   ASVLTA-------GEMAHPINR-RLKV-MVGVINEIQRRAAEACKV--VPTKADTIAAAV   186
CatmPrx     ASLLLDPVA-------------------------------------G--KVSCADILAQAA    97
SsPrx03     GSILLNGTG-TNLAEKDNPDVK-HLR--GFDALERVKATVEASCPR--KISCADTLTLMT   138
KnitPrx     ASVLLNSTA-SISGEKDAFQSM-TLA--GFDVIDDIKAHIETVCPS--TVSCADIVALAS   135
HRPC        ASILLDNTT-SFRTEKDAFGNANSAR--GFPVIDRMKAAVESACPR--TVSCADLLTIAA   135
PNP         ASVLLDDTS-NFTGEKTAGPNANSIR--GFEVIDTIKSQVESLCPG--VVSCADILAVAA   127
```

```
PpPRX1      QIAVKKLSGG--KITWKVP------LGRRDGLVSS-AADVA-GKLPAPTANVATLKSIFA   178
PpPRX38     QISVRLLSDG--TITYPVA------LGRRDGLVSN-ALLVT-GRLPPPTASATTLKLLFK   182
PpPRX4      RDSVVLAG----GYGWRVP------AGRRDGKVSL-AEEATQMNLPAPNATVSQLIRMFG   180
PpPRX49     RDAVVLAG----GYGWHVP------AGRRDGTVSI-MEEA--LNLPAPSMTVSQLIDVFG   178
PpPRX50     RDAVVLAG----GYGWHVP------AGRRDGTVSI-MEEA--LNLPAPSMTVSQLIDVFG   178
PpPRX25     RDSVILAN-------WRLPMERARWASRRDDFPHH-GGNGC-EPRPSEHDPVSTR-SKVP   185
PpPRX46     RDVVILAG----GFGWNVP------GGRRDGTVSK-VEDVNLTNLPSPRANATNLIQLFE   192
PpPRX6      RDTVRITG----GSSWKVY------GGRRDGLISN-AVEVA-QNLPPSTAKVPELVATFA   176
PpPRX23     RDTVILTK----GVGWEVP------AGRMDGRISL-STEPL-QELPPSTFTSQQLISIFA   175
PpPRX31     RDAVVLTG----GKGWRVI------AGRRDGRISN-KIEPE-QNIPTAFASVNELVSTFA   176
PpPRX36     RDVVILTG----GCDWRVL------AGRRDGLVSN-STEVP-KNILAPDKKVSDLLQAFQ   161
PpPRX5      RDSVLLTG----GEGWDVS------GGRRDGFTSS-SADPP-LELPLQTMTVPELLANFA   178
PpPRX33     RDSVTLTG----GEGWDVS------GGRRDSLVSS-YVDPP-LGLPLQTDTVSELLANFA   187
PpPRX12     RDSVLLTK----GKGWTVY------GGRRDGTVSN-AADPP-NNLPVPTMTPTQMIPLFA   188
PpPRX17     RDSVLLIK----GKGWTVY------GGRKHGTVSN-SADPP-INLPVETQTSSQMIPIFV   187
PpPRX7      RDAVIMTG----GLGWPVE------AGRLDGRSSD-ASRAN-AEIPDPSFNVAQLIDSFA   184
PpPRX26     RDAVVLTG----GLGWAVE------AGRLDGRVSD-AGRAF-AEIPDPSFSSAQLAAVFA   184
PpPRX14     RDAVELMG----GLGWRVR------AGRYDGRVSS-AARAL-AEIPDPRYTVEEITALFA   185
PpPRX8      RDSIELTG----GKRWEVP------AGRRDGNVSI-NAEAE-AMLPSPQLNVQQLTDSFT   177
PpPRX11     RDAVLLTG----GQHFDMP------TGRLDGMVST-ADNAN-NNLVSTRSSATELTRKFL   181
PpPRX19     RDAIVLTG----GRHFEMP------TGRLDGMVSS-TASAD-ANLVSTESSARELTQKFL   178
PpPRX24     RDSAVLAGLNFQGLPLTMA------TGRWDGRVSS-RNAAE-AALPSSKSNVQQLTAQFS   191
PpPRX35     RDGAVLAGLNFEGRPLTMA------TGRWDGRVSS-MSAAA-AALPSSKSNVQQLTAQFG   190
PpPRX9      RDSVALSG----GPYWEVQ------LGRRDSLTAS-KTDAE-NSIPQPTFTVTQLVASFN   190
PpPRX15     RDAVFLSG----GPFWDVE------LGRRDALTTS-SQAAV-NSIPSPRFNVPQLIKSFN   186
PpPRX37     RDSVVQTG----GPHYDVL------LGRRDSIIAN-YTGAN-AVLPSPKFNVTTLTKKFL   185
PpPRX16     RDSAVEVGL---SERYPVF------VGRLDSLNAS-RDEAN-LRLPSPRANYSELKKNFE   191
PpPRX48     RDSAVQVGL---TDTYPVY------FGRRDSLTAS-IDEAN-LRLPTPNSNYSVLKANFE   181
PpPRX47     RDSAVEVGL---VPEYPVP------FGRRDSLRAAPIAEVN-LRLPGPDFDISTLKASFA   179
PpPRX2      RDSVEILG----GATYGAE------TGRFDGA----APAAS-VNIPSPNSAVAEATPFFT   181
PpPRX10     RDAVRLSG----GPNFAMP------TGRRDGRVSR-ADN---VNLPGPTVSVADATRIFN   178
PpPRX18     RDVIALSG----GPKFAMP------TGRRDGRVSK-ASN---VNLPGPSLSVADATRAFT   178
PpPRX30     RDGVRLTG----GFFYRVP------AGRRDGYDSI-AAEAT-KNLPDPRMNVDQLTLNFK   192
PpPRX3      RDAVVLAG----GPTWHVE------LGRRDGRISR-ADQAG-SQLPSSQSTAESLITQFA   194
PpPRX32     RDAVVQAG----GPTWSVE------LGRRDGRVSR-ADQAG-SMLPSSQSTAESLIVQFA   194
PpPRX13     AEAVEQAG----GPHVDLA------YGRRDGLESF-APAAK-TNLPAGTLQVAGLLENFQ   229
PpPRX29     AEAVEQAG----GPHIDLA------FGRRDGLDSF-ALAAK-TNLPGSTLKVVGLVENFA   220
PpPRX51     VEAVEQAG----GPHIDLA------YGRRDGLDSF-GQAAK-TNLPGSTLQVEGLRENFA   192
PpPRX28     VECVEQSG----GPHIDLA------YGRRDGLESF-AAAAA-TYMPGGFLRVQGLIESFQ   213
PpPRX22     AKAVELAG----GPILVTE------TGRRDGVVSY-LAGAT-ASMPLSTQKIPDLEAMFV   198
PpPRX44     AKSVELAG----GPILQTE------TGRRDGVVSY-LAGAT-ASMPLSIQKIDDLLAMFV   195
PpPRX20     RDAVSYTG----GPEIPIP------LGRKDATTAS-SENAD-DQLPPASSTVSTMLQVFS   174
PpPRX21     RDAVAYNG----GPEIQIP------LGRKDADFSS-ATEAE-AKLPPATSNVDRVLNVFA   174
PpPRX43     RDAVAFSG----GPEIQIP------LGRKDADFSS-ASEAD-AKLPPSTSSVDTILSVFA   180
PpPRX40     RDAVAFNG----GPDIKIP------LGRKDAVSSS-ATEAD-AKLPPATSSIDRVFNVFG   179
PpPRX42     RDAVAFNG----GPDIQIP------LGRKDADSSN-AGEAD-SKLPPATSSIDRVFNVFG   180
PpPRX34     AAGAQVLG----GPAIKLK------TGRKDSRVSL-KSVAD-TGIPTPQSNVSFVLDYFS   175
PpPRX39     AAAVQVLG----GPYIHVK------TGRKDTRNSM-KSSAD--TIPRPQDGVTKVLTFYK   176
PpPRX41     RDAYVLASK---RPAYPVL------LGRLDGMVSL-VKDAD--SLPGARTPINTIVSVFK   214

CbraPrx01   IASAEFYGLSC-ARYCPFK------LGRRDVHPSS-KGDDP-GTLPSVTDSVSIVLEKMG   237
CatmPrx     VEAVHRVG----GPKISLR------MGRLDGNVSL-ADDVL-RFLPSPFFNTDQLIATFE   145
SsPrx03     KFAVIKSG----GPEIDFN------LGRLDGFSSN-LELAK-LLTPRHFSTATALLNIFF   186
KnitPrx     TESIFQSG----GPNISMP------LGRVDGRVSS-KAAAL-AAMPNVTMNVTVLNASFA   183
HRPC        QQSVTLAG----GPSWRVP------LGRRDSLQAF-LDLAN-ANLPAPFFTLPQLKDSFR   183
PNP         RDSVVALG----GASWNVL------LGRRDSTTAS-LSSAN-SDLPAPFFNLSGLISAFS   175
```

```
                                P      6
PpPRX1     GVGLTT-EEMVVLS-GAHSVGVASCRAV-QNRLTT--------PPDATLDPTYAQALQRQ   227
PpPRX38    AVGLST-EDMVVLS-GAHSIGKARCSFF-RNRLTT--------PSDANMDPDYAESLKRQ   231
PpPRX4     AKGLSA-SEMVVLS-GAHTIGRAPCVTF-DDRVQT-------SPVDPTLAPNFAASLKRQ   230
PpPRX49    RKGLSP-SQMVVLS-GAHTIGKAPCVTF-DDRVQT-------TPVDPTLAPSFATFLKGQ   228
PpPRX50    RKGLSP-SQMVVLS-GAHTIGKAPCVTF-DDRVQT-------TPVDPTLAPSFATFLKGQ   228
PpPRX25    GEGLER-SLMVVLS-GFHTIGKAPCLTF-DDRVQT-------NPVDPTLAPSFAASLKKQ   235
PpPRX46    SKGMTL-AQMVVLS-GAHTIGKATCITF-DNRLHSDIPEP---YADP-YSPSFKLYLKSQ   245
PpPRX6     QKGLTP-QQMVDLS-GSHTLGVTHCVHL-RDRIFT--------PIDPTMPKSLLKQLQRV   225
PpPRX23    GKGLTA-KQMVDLS-GSHTLGITHCLHL-RDRIFT--------TIDPTIPKNLLRQLQRK   224
PpPRX31    QQGLNT-EDMVVLS-GAHTIGVTHCNHI-SDRIYN--------PVDKTMPKDLLKSLQKS   225
PpPRX36    KKGFNA-AQMVTLT-GAHTIGRASWFAF-DVRIHNFSGDQ--SKVDPSLPPLFASILKKK   216
PpPRX5     AKNLNA-AHMVALS-GSHSIGVAHCQFI-VDRLYNYPNSAT--GSDPSLPADLLEFLKTQ   233
PpPRX33    EKNLNA-AHMVALS-GGHSIGIAHCQYV-TDRLYDYPSSDT--GSDPTLPSDMQATLKTE   242
PpPRX12    GKGLSA-DDLVALS-GSHTIGIAHCIFV-NPRIYG-------NNTDPTIPADFLASLKSQ   238
PpPRX17    SKGLSA-DDLVALS-GGHTIGIAHCTFV-SPRIYG-------NNTDPKIPADFLASLKRQ   237
PpPRX7     RKGLTR-SDMIVLS-GAHTIGRANCKSV-ATRLYP--------VQDPRLSEPLAAELKSG   233
PpPRX26    RKGLTT-SDMIVLS-GAHSIGRAHCDSV-KTRLYP--------VQDPNLREPLAAELRSG   233
PpPRX14    RKGLSK-SDMIVLS-GAHTIGRAHCASV-TPRLYP--------VQDPQMSQAMAAFLRTA   234
PpPRX8     RKGLSQ-SDMITLS-GAHTIGRIHCSTV-VARLYP--------ETDPSLDEDLAVQLKTL   226
PpPRX11    EQGLGQ-DDMITLS-GAHTVGKTTCGQI-TSRLYNFPGTT--NGVDPTLDFDYALHLQQL   236
PpPRX19    AQGLGQ-DEMITLS-GAHTIGRTTCAQV-TPRLYNFPGSP--NGVDPTLDFDYALHLKQV   233
PpPRX24    NKGLSQ-DEMVTLS-GAHSIGVAHCSNF-MDRLYDFPGSP--NGVDPTLDPDYAAELQAK   246
PpPRX35    AKGLSQ-DEMVTLS-GAHTIGKAHCVNF-MDRLYDFPGSA--TGVDPTLDANYAAELQTQ   245
PpPRX9     AVGLNE-KDVVALS-GSHSFGKARCTSF-QNRLGNQASGSQSPGSDPFLESSYLAKLQTL   247
PpPRX15    AVGLDK-KDVVALS-GSHTIGIARCASF-QARLYNQGNSG---RPDSSLEKHYLAELQNR   240
PpPRX37    DVGLTS-EDMVTLS-GAHTIGKTHCTSI-TTRLYNQSGTT---KPDPAIPAEMLRKLQTK   239
PpPRX16    FQGLNE-VDLIALS-GAHTIGKVRCQIV---RLF---------LNDTDTNAEFRSNLAKE   237
PpPRX48    FQGLDE-TDLIALS-GAHTIGRVRCIVI-TVSNS---------STDPNINAAFRDTLIKA   229
PpPRX47    NQSLDE-RDLVALS-GAHTIGRVRCQFV---RLF---------LNDPGTNADFKKELARL   225
PpPRX2     NLGLTQ-DDMVNLL-GAHTVGVSQCQFF-VDRLYNFQGTG---LPDPSLDATYLAVLQSR   235
PpPRX10    AQGLTR-NDMVTLL-GAHSVGITHCSFF-HERLWNFEGTG---SADPSMDPNLVMRLKAI   232
PpPRX18    AQGMTQ-NDMVTLL-GAHTVGITHCSFF-DDRLWNFQGTG---RADPSMDANLVKQLKSV   232
PpPRX30    NQGLTR-DEMVILS-GAHTIGDVACHHI-DNRLYTYPGNN---GVVPSLPRAFVKKLKGI   246
PpPRX3     ALGLTP-RDMATLS-GAHTFGRVHCAQV-ARRFFGFNSTT---GYDPLLSDTYATKLRTM   248
PpPRX32    AMGLTP-RDMATLS-GAHTFGRVHCAQV-ARRFFGFNSTT---GYDPLLSETYAIKLRSM   248
PpPRX13    NVGLNL-TDVVVLS-GGHTIGQARCSSF-ADRFTP-------GVKNPFPDVSFGENLYTY   279
PpPRX29    NVGLNK-TDMVVLS-GGHTIGQARCSTF-ADRFAP-------GVKNPFPDTKFGEALQTY   270
PpPRX51    NVGLNL-TDMVVLS-GGHTIGQARCSSF-ADRFAP-------AAKNPFPDTIFGQELNAY   242
PpPRX28    MAGLDE-VDLVALS-GAHTLGQARCSEFIQERFIS-------PGSNSFRDSDYGLALQSY   264
PpPRX22    QAGLDI-NDLVILS-GAHTIGEVHCSNF-ADRFDP-------AANSPFGDVSFGQELLAF   248
PpPRX44    QAGLDL-TDLVILS-GAHTIGEVHCTNF-ADRFDP-------AANSPFPDPSFGEQLRAY   245
PpPRX20    RYGMTA-AETVGIL-GAHTLGIGHCVNV-VDRLYP--------TRDPALSTGLYLQLRVL   223
PpPRX21    PFGMSI-AESVAIL-GAHTLGVGHCKNI-QDRLQL-------NSPTAPNSVVYRTQLRAA   224
PpPRX43    PFGMSL-AESVASL-GAHTLGGGHCKNI-QDRLRF-------NSPTAPTSLLYRTQLRAA   230
PpPRX40    AFGMTH-EESVAIL-GAHTIGVGHCKSI-QDRLQS-------NSPTAPNSLVFRTQLTAA   229
PpPRX42    PFGMTP-EEIVAIL-GAHSIGVGHCKNI-QDRLQS-------NSPTAPNSLVFRTQLMAA   230
PpPRX34    KMGINT-EETVALL-GAHTIGRAHCVSF-EERIYP--------TVDPKMDPVFASMLKYR   224
PpPRX39    NIGINP-REAVALM-GAHTIGRAHCTSF-IERIFP--------KVDPKMDPVFAEKLKRR   225
PpPRX41    DAGFTT-EEAVILS-GAHTIGEAKCKFF-NDRLHNFLNTK---KPDRTMDPALVEQLKKI   268

CbraPrx01  SLGFSP-KEVALYSLGSHSIGQASCFLF-EDRLNKRQCRL---PRDERLDNGRACDLVKV   292
CatmPrx    AVGLGA-DDVVPLS-GGHTFGETHCVAV-RPRLDA-------PGGDPALTPSHAAFLEQT   195
SsPrx03    LLGLSQ-VDLVALS-GAHTIGVAHCESV-RTRIYP--------TIDTKYEPQFGISVRKS   235
KnitPrx    AVGLTL-GDMVILS-GAHTFGKAHCSNV-VDRLLP---------VDPTLDPALAQNLTQQ   231
HRPC       NVGLNRSSDLVALS-GGHTFGKNQCRFI-MDRLYNFSNTG---LPDPTLNTTYLQTLRGL   238
PNP        NKGFTT-KELVTLS-GAHTIGQAQCTAF-RTRIYN----------ESNIDPTYAKSLQAN   222
```

```
              7                                              P
PpPRX1        CPAGSP----------NNVNLDVTTPTRLDEVYFKNLQARKGLLTSDQVLHEDPET---K    274
PpPRX38       CPADKP---------NNLVDLDVTTPTNLDSEYYKNLQVNKGLLTSDQNLQSDPET---Q    279
PpPRX4        CPYPGI--------GSTSVNMDS-TTRRFDSQYYKDIIRGRGLLTSDQGLLYDSRT---K    278
PpPRX49       CPYAAI--------QSTSVDMDS-TAHTFDSQYFKDIIAGRGLLTSDQSLLYDSRT---S    276
PpPRX50       CPYAAI--------QSTSVDMDS-TAHTFDSQYFKDIIAGRGLLTSDQSLLYDSRT---S    276
PpPRX25       CLYAQI--------TSTKVALDS-TPRRFDTQYFKDIIQGRGVLISDQELLYDSRT---V    283
PpPRX46       CPNPNM---------FVRVNLDS-TPEKFDGRYFHDLVHHRGLLTSDQTLMSDSRT---R    292
PpPRX6        CPKITS---------PTPLVIDRLTPHKFDTQYYQNIASGQGLMTSDQDLFNDDST---R    273
PpPRX23       CPSNTS---------LTPLQIDRYTGNKFDTQYFRNIVRGRGLMTSDQDLFRDPAT---K    272
PpPRX31       CPKASS---------PTSLVMDRKSVHKFDTEYFRNIRAGYGLMTSDQGLYREDFT---R    273
PpPRX36       CPSANL---------TKWVNLEVITPRRFDTQYYKNLIHKIGLLTSDMSMVADSHT---Q    264
PpPRX5        CPDSAA---------TPEINIDEVSPGTFDSQYFDNIIRNRGVIASDQHLMDHTST---Q    281
PpPRX33       CPNAAA---------TPELNVDEVTPDTFDSQYFNNIVKGRGLLASDQRLMDDKAT---S    290
PpPRX12       CPADSVTTN---PPVGAPINLDRVSPTKFDSQYFQNIIDRKGLLTSDQSLLDDSRT---R    292
PpPRX17       CPADSVTTN---PPIGAPIDLDLVSPTKFDSQYFQNIIQRKGLLTSDQSLLDDSRS---R    291
PpPRX7        CPQQGG---------SATFNLDS-TPDRFDNNYYANVVNGRGIMNSDQVLFDDPST---R    280
PpPRX26       CPQQGG---------SATFSLDS-TPNQFDNAYYIDVVNGRGIMRSDQALFDDPST---R    280
PpPRX14       CPPQGG--------SAATFSLDSTTPYRFDNMYYTNLIANRGLLHSDQALINDMST---R    283
PpPRX8        CPQVGG-------SSSSTFNLDPTTPELFDNMYYSNLFSGKGVLQSDQILFESWST---K    276
PpPRX11       CPQNGN--------PNDPVPLDPVSPNTFDNMYYTNGVTGRVLFPSDNVLFADHQT---Q    285
PpPRX19       CPQGGN--------PNSVVQLDPVSPNTFDNMYYTNGVTGRVLFASDIALFADHQT---E    282
PpPRX24       CPRGNP-------NPNTVVNMDPQTPFVIDNNFYSNGFAGKVLFSSDMALFNDFET---Q    296
PpPRX35       CPRGNP-------NQNTVVDLDPATPFVMDNNYYRNGFAGKVLFGSDMALFHDFET---Q    295
PpPRX9        CPSNGD--------GNTTVNLDHFTPVHFDNQYYKNLQAAKGLLNSDAVLHTTNGQ--SN    297
PpPRX15       CPQSGD--------GNQTAFLDPCTPTTFDNQYYKDLQAGRGLLFSDEVLETTSGT--TL    290
PpPRX37       CPNDPT-------DLKTTLVLDDETPEVFDNQYFKNLLNKRGILYSDQILADTEGF--NL    290
PpPRX16       CPAGGD--------DFKLQNLDLKTPEKFDNNYFKNLRRGEGIIRSDQTLWSTPGI--NQ    287
PpPRX48       CDTANG------TIDPPLQNLDVKTPDKFDNNYFKNLRRGEGVLTSDQTLQSTPGP--NV    281
PpPRX47       CAPTVD--------AFTLQNLDLKTPDKFDNNYYKNLRRGEGIIRSDQVLWSSEGT--HQ    275
PpPRX2        CPNVAG--------DVTTVALDQGSESSFDTGYFTNIQASKGVLRIDQEIANDAST---S    284
PpPRX10       CPQQGV-------GLGSPVNLDQATPNIMDNTFYNQLIARKGILQLDQRVATDRTT---T    282
PpPRX18       CPQRGV-------GLGRPVNLDQGTPNIVDKVFYSQLLAKKGILQLDQRLATDRAT---S    282
PpPRX30       CPRPNL--------FDITVDMDQVTPIRFDSQYYKNLASKTSVLSSDQVLYDDVRT---R    295
PpPRX3        CPQPVD--------GTSRIPTEPITPDQFDEHYYTAVLQDRGILTSDSSLLVNAKT---G    297
PpPRX32       CPQPVD--------NTARIPTEPITPDQFDENYYTSVLESRGILTSDSSLLINVKT---G    297
PpPRX13       CVEGNT------IGLDRRMSLDTNSTTVFDNGYFRSLVAGRGILTSDNILFTDPRT---K    330
PpPRX29       CTDGNT------AGLDRRMTLDANSTTVFDNGYFRSIVAGRGILTSDHVLFTDPST---K    321
PpPRX51       CVEGNR------LGIDRRMTLDANSTTIFDNGYFQSIVAGRGILTTDNVLFTDNRT---K    293
PpPRX28       CAEGKN------LGLDRKVTLDSNTSTIFDNGYFQTLVDGRGVLTSDNDLTLDNRT---A    315
PpPRX22       CTRNGAGDI---ATLNLKTFMDLQTPNSFDISYYVNLIIGRGVMTSDQVLFNDLRT---Q    302
PpPRX44       CTRGGTGDM---ATLNLRTFIDLQSPNSFDISYFVNLIVGRGVMTSDQALFNDQRT---Q    299
PpPRX20       CPTKEP--------LNLTILPNDLSVYSFDNRYFKDVLGGRGLFRADANLVGDART---K    272
PpPRX21       CAVNVF---------DIAILNNDASQFTFDNQYFQDIQNGRGLFTVDDQLSTDPRT---A    272
PpPRX43       CVVNVF---------DIAILNNDASQFTFDNQYFKDIQNGRGLFTVDNLLSTDPRT---A    278
PpPRX40       CAVNVF---------NIAVLTNDATQFTFDNQYFKDIQNGRGLFTVDNLLSIDPRT---A    277
PpPRX42       CAVNVF---------DIAVVNNDATQFTFDNQYFQDIQNGRGLFTVDHLLSTDPRT---A    278
PpPRX34       CPQQKTGAE---PVHFTYFRNDEQSPMAFDNHYYVNLMANQGLLHIDSEIAWDSRT---K    278
PpPRX39       CPAKPT------SVHFTYFRNDEPSPMAFDNNYFKNLVTKQGLMGIDSALYWDGRT---Q    276
PpPRX41       CPNMTA-------NLESPAFLDQGTRRVFDKSYFVQVTRQRGVLQSDQNLFANAAT---K    318

CbraPrx01     CAA----------KRNQLVPFDYITPTKLDTNYLKLVLQRQGLLRIDQDFGDDART---A    339
CatmPrx       FPDPTCGGGNPFAPGGPAVNLDNITSDRFDGGYFKGIRNGRTAMRSDAALL-EGQL---K    251
SsPrx03       CPIVDPSQNPPLLDPNATLKLDQ-TSTVFDNSYFQGLRSNRGLMTSDNNLNADPRL---N    291
KnitPrx       CNPRVN-------DPTITVPLDVATNATFDNQYYKNLLQGRGVLASDEVLAHDNRT---V    281
HRPC          CPLNGN--------LSALVDFDLRTPTIFDNKYYVNLEEQKGLIQSDQELFSSPNATDTI    290
PNP           CPSVGG--------DTNLSPFDVTTPNKFDNAYYINLRNKKGLLHSDQQLFNGVST---D    271
```

```
                                                                      8
PpPRX1      PMVAKHT---SQGVF-----NEAFKNAMRKMSDIGVLTG---SAGEIRANCHRFNA----   319
PpPRX38     PMVSDNA---EPGTF-----RTKFADAIRRMSNIGVLTG---SAGEIRLNCRRFN-----   323
PpPRX4      RDVHAN----KGSAF-----YRNFAQAMVAMSRIEVLTG---RSGEIRRQVGEVNKY---   323
PpPRX49     GGVYAN----NGAAF-----YRNFAKAMVKMSQIEVLTG---LDGEIRRQFDQVNSH---   321
PpPRX50     GGVYAN----NGAAF-----YRNFAKAMVKMSQIEVLTG---LDGEIRRQFDQVNSH---   321
PpPRX25     GVVRAN----KGSAF-----YRNFGEAMVAMSELGVLTD---GSGEIRRQIDQVNT----   327
PpPRX46     HCVYKNR---DDGVF-----KKNFAEAMVAMSKIGVLTG---KDGEIRRRMEVVNSK---   338
PpPRX6      RFVVKNL---KHGNF-----IHRFGKAMIAMTNIEPTIA---PDGEIRRRCQFLN-----   317
PpPRX23     PFVEANL---KRATF-----DKNFAEAMVAMTSIEVKIG---HEGEIRKHCQFVN-----   316
PpPRX31     PIVDANL---NQRAF-----VNRFAEAMFKLQFIQPLEA---PDGEIRRRCQCRN-----   317
PpPRX36     EQVYMNT---NWQKF-----SSNFADAMVDLSKLDVLTV---QSGEIRLKCRFVN-----   308
PpPRX5      GEVAAN----NGPAF-----GGNFGRAMVVMARFNVLTG---SAGQIRTNCRQVN-----   324
PpPRX33     DAVLAN----NGPDF-----GGNFGRAMVVMARYNVLTG---NAGQIRTNCRQVN-----   333
PpPRX12     GAVYKN----SGNFF-----NSEFGRAMQAMAGIGVLTG---NEGQIRTNCRAVNP----   336
PpPRX17     NAVYKN----NGRFF-----NSEFGRAMQAMARVGVLTG---NQGQIRKNCRALNP----   335
PpPRX7      PETTFNAV--GSAPW-----AFRFSQIMLKMGTIDVKTG---PQGEIRRNCRSVN-----   325
PpPRX26     TETMFNSL--GAAPW-----AFRFGQIMVKMGQVGVKTG---PDGEIRRNCRFVNTPI--   328
PpPRX14     GETIFNSF--AAGPW-----AFQFSRVMIEMGNIQVKSG---PDGEIRRHCRFIN-----   328
PpPRX8      LPTMFNVL--STTSF-----TSSFADSMLTMSQIEVKTG---SEGEIRRNCRAVNPVVEA   326
PpPRX11     FASNLNSQ--NGQFW-----QMKFANALVRMASNKVKLGVPNRNGEIRKNCRFTNAATGS   338
PpPRX19     FASNLNSE--NAELW-----QIKFSNALIHMASNKIKFGRPDEEGEIRQNCRLTNARFAA   335
PpPRX24     FTSDLNVV--NGITW-----NQKFGNALAQMAAIDIKDD---FDGEVRLNCRRIN-----   341
PpPRX35     FTSDLNVV--NGVSW-----NQKFGNALAQMASIEVKDS---TVGEIRLNCRRVN-----   340
PpPRX9      QLVEIYAN--DERVF-----FKDFAQSVLKMGSIKVMTG---NKGEVRRNCRLPNTIRA-   346
PpPRX15     KLVELYAT--DQTAF-----FTDFVSSMLKMASIHVKAD---SEGEIRRNCRIPNSVNAK   340
PpPRX37     DLVNLYAN--DQNAF-----FDAFVKSMTRMGNISPLMG---TSGEIRKRCDRVN-----   335
PpPRX16     AIVWDFAR--NQKTF-----FRQFAFSTIKMGNIRPPAG---TKGEIRENCRAVNSAP--   335
PpPRX48     GIVKDFAK--NKENF-----FTQYGLSSIKMGYIRPLTG---DQGEIRKNCRAVNSAPSS   331
PpPRX47     KITKDFAE--NQENF-----FRQFIESSIKMGKIKPPPG---SPSEIRLNCHQANPRP--   323
PpPRX2      GRVNTLAA--SPSTF-----GTDFATSMIAMGRIAVL-----TSGSVRSDCETA------   326
PpPRX10     ARVNVLAS--PRSTF-----TAAFAASLIRLGNVRVIEG---SGGEIRKICSRIN-----   327
PpPRX18     QRTRTLAG--PTSPF-----TKDFVAAIIKLGNVKVLEG---TKGEIRKICSRIN-----   327
PpPRX30     PLVRVLE---SKLAF-----LSKFGPAMVRMGNINVLTG---NQGEVRLNCRRKNSPGSA   344
PpPRX3      RYVKEYAQ--NRTVF-----FERFAAAMLKMGRFGVKLG---TEGEIRRVCSAVN-----   342
PpPRX32     RYVTEYAN--NRSVF-----FERFTAAMLKMGRVGVKLG---SEGEIRRVCSVVN-----   342
PpPRX13     PLVTQFAE--NQDAF-----FTAFKESMAKMGRIVVLTG---TQGQIRKQCWVRNPIDAN   380
PpPRX29     PLVTLFAA--NQDAF-----FAAFKESMAKMGRIGVLTG---TQGQIRKQCWVRNPVDIT   371
PpPRX51     SLVTTFAQ--DQTVF-----FDAFKELMAKMGRIGVLTG---TQGQIRKQCWVRNPIDPA   343
PpPRX28     PLVQLYAS--DQNAF-----FTAFAASMRKMSKIGILTG---TQGQVRKKCYVRNSVDVV   365
PpPRX22     PMVREFAA--NRTLF-----FESFQASMLKMGRLHVLTG---TNGVIRKQCGVYP-----   347
PpPRX44     PLVRAFAG--NRTLF-----FESFQASMLKMGRLHVLTG---TSGVIRRQCGVYP-----   344
PpPRX20     PLVAKFAS--DQSLF-----FKTFASAYVKLVSAQVLTG---SRGEVRTNCRRVNAQD--   320
PpPRX21     PIVTLYAS--NQGAF-----FSAFQSAYVKLTS-RAMTG---NQGSVRSTCTL-------   314
PpPRX43     PIVSLYAT--NEAAL-----FAAFQSAYVTLTS-RAMTG---TQGSVRSTCHL-------   320
PpPRX40     PIVNTYAA--NKGAF-----FAAFQSAYVKLTS-RALKG---NQGSVRSTCLH-------   319
PpPRX42     PIVNTYAS--NEGAF-----FASFASAYVKLTS-RAVTG---NRGSVRSTCQL-------   320
PpPRX34     LFVVEYAK--DNALW-----HKNFATAFTKLSEHNPLTG---TQGEVRKHCSYTL-----   323
PpPRX39     KYVIEFSQ--NEAAW-----REVFTVAFKKLSEYKVLTG---RQGEIRKRCMYVN-----   321
PpPRX41     QFVTGLASG-TEENF-----LSKFEKAMAKLGNLGATTN---YKGNIRRVCSVLN-----   364

CbraPrx01   GVVRQYASGKGERSF-----HGDYLQVLVKSLSIGI-NG---PKSVAPRKYRLTT-----   385
CatmPrx     DQVLLYAR--DRAAF-----FTNFTRSLRRMSKLGVKLG---EEGEIRTNCRRVN-----   296
SsPrx03     SQVAKFAS--NQSAF-----FNQFRLSMKKLSEINLR-----SVGNVRRKCYKLN-----   334
KnitPrx     KLVQRMAT--NERAF-----FKNFSHAWVRMSMIGVKTG---QEGEIRRNCGVVNRR---   328
HRPC        PLVRSFAN--STQTF-----FNAFVEAMDRMGNITPLTG---TQGQIRLNCRVVNSNSLL   341
PNP         SQVTAYSN--NAATF-----NTDFGNAMIKMGNLSPLTG---TSGQIRTNCRKTN-----   316
```

```
PpPRX1      ---------------------------------------
PpPRX38     ---------------------------------------
PpPRX4      ---------------------------------------
PpPRX49     ---------------------------------------
PpPRX50     ---------------------------------------
PpPRX25     ---------------------------------------
PpPRX46     ---------------------------------------
PpPRX6      ---------------------------------------
PpPRX23     ---------------------------------------
PpPRX31     ---------------------------------------
PpPRX36     ---------------------------------------
PpPRX5      ---------------------------------------
PpPRX33     ---------------------------------------
PpPRX12     ---------------------------------------
PpPRX17     ---------------------------------------
PpPRX7      ---------------------------------------
PpPRX26     ---------------------------------------
PpPRX14     ---------------------------------------
PpPRX8      P---SPL--------------------------------   330
PpPRX11     S---TNN--------------------------------   342
PpPRX19     T---ANMSRSAHN-----------------------HGA   348
PpPRX24     ---------------------------------------
PpPRX35     ---------------------------------------
PpPRX9      ---------------------------------------
PpPRX15     G------------------------------GT------   343
PpPRX37     ------------------------L--------------   336
PpPRX16     -----LVASE-----------------------------   339
PpPRX48     -----LVAYQ-----------------------------   336
PpPRX47     -LIEQVVAVE-----------------------------   332
PpPRX2      ---------------------------------------
PpPRX10     ---------------------------------------
PpPRX18     ---------------------------------------
PpPRX30     ---------------------------------------
PpPRX3      ---------------------------------------
PpPRX32     ---------------------------------------
PpPRX13     L---RPEANMDFAPQSTKFCRQKP-----CNPTCSDPAS   411
PpPRX29     T---TPDANMDFAPVSSEFCKPAS-----CDATCASPTS   402
PpPRX51     T---TPDANMEFAPASLEFCTPNP-----CPATCSSNTA   374
PpPRX28     K---SPNSNTEFSPISPTICKPAPQVDQKCDGT------   395
PpPRX22     ---------------------------------------
PpPRX44     ---------------------------------------
PpPRX20     ---------------------------------------
PpPRX21     ---------------------------------------
PpPRX43     ---------------------------------------
PpPRX40     ---------------------------------------
PpPRX42     ---------------------------------------
PpPRX34     ---------------------------------------
PpPRX39     ---------------------------------------
PpPRX41     ---------------------------------------

CbraPrx01   ---------------------------------------
CatmPrx     ---------------------------------------
SsPrx03     ---------------------------------------
KnitPrx     ---------------------------------------
HRPC        H---DMVEVVDFVSSM-----------------------   353
PNP         ---------------------------------------
```
